# SimUrine: A Novel, Fully Defined Artificial Urinary Media for Enhanced Microbiological Research of Urinary Bacteria

**DOI:** 10.1101/2025.08.01.668053

**Authors:** Pablo Gallardo Molina, Brian I Choi, Michael Vanek, Mohammad Haneef Khan, Marjon de Vos, Alan J. Wolfe

**Affiliations:** Department of Microbiology and Immunology, Loyola University Chicago, Maywood, IL USA; GELIFES-University of Groningen, Groningen, The Netherlands

**Keywords:** Artificial urine, Urobiome, Urinary tract infection, Microbiome research, Bacterial cultivation, Microbial ecology

## Abstract

Urinary tract infections represent one of the most prevalent bacterial diseases, yet current diagnostic and research methodologies are hampered by inadequate culture media that fail to replicate the bladder biochemical environment. Conventional artificial urine formulations contain undefined components, lack essential nutrients, or inadequately support urinary microbiome (urobiome) growth. To address these limitations, we developed SimUrine, a fully defined synthetic urine medium that aims to replicate human bladder chemistry while supporting diverse microbial growth requirements.

SimUrine was systematically developed through iterative optimization of multi-purpose artificial urine, incorporating defined concentrations of carbon sources, vitamins, trace elements, and amino acids within physiologically relevant ranges. The modular design enables component substitution without complete reformulation, facilitating customization for culturomics, antimicrobial susceptibility testing, and microbial ecology studies, while reducing batch-to-batch variability associated with authentic urine.

Performance evaluation demonstrated SimUrine’s capability to support growth of fastidious urobiome members, including *Lactobacillus* species, *Aerococcus urinae*, and *Corynebacterium riegelii*, which fail to proliferate in conventional minimal media. Physicochemical characterization confirmed that SimUrine formulation exhibits properties within normal human urine ranges for density, conductivity, osmolarity, and viscosity, ensuring physiological relevance.

Clinical applications revealed reduced antibiotic susceptibility compared to standard media, suggesting more accurate representation of *in vivo* conditions. Co-culture experiments using *Escherichia coli* and *Enterococcus faecalis* demonstrated previously unobserved microbial interactions, highlighting SimUrine’s utility for investigating urobiome dynamics.

SimUrine represents a significant advancement in urobiome research methodology, providing a standardized, reproducible platform for investigating urobiome under physiologically relevant conditions, potentially improving fundamental understanding and clinical diagnostic approaches.

**IMPORTANCE:** Urinary tract infections affect millions globally, yet current research and diagnostic methods rely on inadequate culture media that fail to replicate the bladder’s unique biochemical environment. This fundamental limitation has hindered accurate UTI research and potentially compromised clinical treatment decisions. SimUrine addresses this critical gap as the first fully defined synthetic urine medium that mimics human bladder chemistry while supporting growth of diverse urinary microbes. The breakthrough enables cultivation of urobiome organisms in a minimal medium that resembles natural conditions, revealing novel microbial interactions that influence urinary health. Crucially, SimUrine demonstrates different antimicrobial susceptibility patterns compared to standard clinical media, suggesting current testing protocols may inaccurately predict treatment outcomes. This standardized, reproducible platform eliminates the variability of authentic urine samples while maintaining physiological relevance, potentially transforming urobiome research methodology and improving clinical diagnostic accuracy for urinary tract infections worldwide.

## INTRODUCTION

Urinary tract infections (UTIs) are among the most common bacterial diseases worldwide. They are particularly common in females; approximately 50-60% will experience at least one UTI in their lifetime (Al Lawati et al., 2024). The diagnosis of UTIs can be challenging due to the variable presentation of symptoms including pyuria, dysuria, and fever, which are not present in all cases. Symptom inconsistency has provoked debates within the medical community regarding standardized diagnostic criteria (Werneburg et al., 2024). Another significant challenge in UTI diagnosis is the discrepancy between these symptoms and the bacterial counts enumerated from standard urine culture technique – a methodology that often produces false negative results, particularly overlooking fastidious and anaerobic bacteria (Hilt et al., 2014). Newer diagnostic techniques with heightened sensitivity have increased the detection rates of bacteria among symptomatic individuals (Haley et al., 2023; Hilt et al., 2014; Moreland et al., 2025a). However, our understanding of these detected bacteria is limited, especially concerning the role of hard-to-cultivate species of the urinary microbiome (urobiome) in the context of UTIs (Moreland et al., 2025b; Price et al., 2016).

These limitations stem partly from the lack of culture media that accurately replicate the bladder’s metabolic environment. Commercially available media, fortified with general nutrients, fail to replicate the nutritional composition and complexity of human urine (Reitzer & Zimmern, 2019). Using authentic urine samples introduces significant donor-to-donor variability, compromising experimental reproducibility (Asscher et al., 1966; Sreedevi et al., 2024). Another strategy is to use pooled urine, which has been theorized to reduce variation. However, a recent report showed that pooled urine is artificially optimized and not average (Hogins et al., 2022). Thus, without a standardized urine medium, comparing experimental results between experiments and different laboratories remains difficult.

Unfortunately, current formulations aiming to replicate bladder-like conditions either contain undefined components, do not encompass the complete range of essential nutrients, or are inadequate to support growth of the broad spectrum of urinary microbes (Brooks & Keevil, 1997; Zandbergen et al., 2021). Salt-based media formulations have been developed to mimic the electrolyte composition of urine; nevertheless, these fail to sustain microbial growth due to their lack of or limited carbon content and absence of other vital nutrients (Chutipongtanate & Thongboonkerd, 2010; Sarigul et al., 2019). To achieve a standardized synthetic urine medium, it is essential to recreate the conditions that members of the urobiome encounter in their natural habitat (Sreedevi et al., 2024), incorporating carbon sources and vital nutrients necessary for the growth of diverse urinary microbes, including species considered to be pathogenic or commensal.

The objective of this study is to develop a fully defined medium that accurately replicates the chemical composition of urine in the human bladder and accommodates various experimental applications, including culturomics, co-evolutionary research, and synthetic communities. To achieve this end, we modified multi-purpose artificial urine (MP-AU) (Sarigul et al., 2019), incorporating defined concentrations of carbon sources, vitamins and salts, with the aim to maintain medium concentrations within the ranges reported for human urine. To determine the efficacy of our formulations, the experimental media were tested for their ability to support the growth of a diverse set of target microbes.

The resultant medium, termed SimUrine, alleviates the inconsistencies associated with using actual urine and ensures more uniform outcomes in urobiome investigations. SimUrine also exhibits physicochemical properties within the normal range of human urine including density, conductivity, refractive index, and viscosity – making for a physiologically relevant *in vitro* artificial medium.

SimUrine’s composition is structured modularly, allowing for the substitution of individual components to accommodate varying research requirements without requiring full medium reconstruction. The modular design also enables facile, prolonged storage and replacement of components upon their expiration date. SimUrine supports a wider range of species than conventional minimal media while sustaining antibiotic susceptibility testing and displaying comparable or enhanced microbial survival rates, including for many fastidious urobiome members.

In summary, SimUrine addresses the significant constraints of existing artificial urine media by providing a fully defined, customizable, nutrient-optimized medium for diverse research applications, enhancing our comprehension of urinary microbial viability and interactions in closely replicated bladder environments.

## METHODOLOGY: EXPERIMENTAL DESIGN AND MATERIALS

### Bacterial Strains

Urinary bacterial isolates used in this study were sourced from strain collections of the Wolfe lab collection at Loyola University Chicago or the de Vos lab at the University of Groningen (**Supplementary Table 1**).

### Reference Media

Our initial experiments utilized NYCIII medium as either a reference or to start cultures (as indicated in each case), due to the fastidious growth requirements of *Lactobacillus* species. However, to ensure simplicity and reproducibility, we conducted subsequent experiments using Brain Heart Infusion (BHI) medium, except for the antibiotic susceptibility testing, for which Mueller-Hinton Broth (MHB) was used to culture reference strains.

### Optimization of MP-AU for Bacterial Growth

To assess bacterial viability in MP-AU (Sarigul et al., 2019), we evaluated *Escherichia coli* (UMB1180) growth and survival in a modified version (mMP-AU), where K_2_C_2_O_4_.H_2_O was replaced by NH_4_C_2_O_4_.H_2_O. For this aim, we first grew the bacteria in NYCIII medium (Geaman et al., 2024) for 48 hours at 37°C in 5% CO_2_ and diluted to OD = 1. The culture was centrifuged, washed three times using PBS 1X and resuspended in either 1 ml of mMP-AU or M9 salts. A 500 µL aliquot of this resuspension was inoculated into 5 ml of mMP-AU or M9 salts and incubated at 37°C for 4 days in 5% CO_2_ with continuous orbital agitation at 200 rpm. Samples were taken every day, and serial dilutions were plated on BHI agar for enumeration of colony-forming units (CFU/mL) to determine survival.

To determine if an enriched version of mMP-AU could support the growth of fastidious species, we tested urinary isolates of *Lactobacillus jensenii* (UBM8651) and *Aerococcus urinae* (UMB5254, aka type strain ATCC 51268) in mMP-AU supplemented with various glucose concentrations. To evaluate additional nutritional requirements, mMP-AU was fortified with an equimolar carbon source (1% each) mix (glucose, galactose, maltose, sucrose, fructose, mannitol, and sorbitol), a vitamin mixture (thiamin 5.0 mg/L, riboflavin 5.0 mg/L, niacin 5mg/L, pantothenic acid 5.0 mg/L, pyridoxine hydrochloride 10.0 mg/L, biotin 2.0 mg/L, and 4-aminobenzoic acid 5.0 mg/L) and casamino acids. Bacterial cultures were evaluated daily for survival, enumerating CFU/mL on Colistin Nalidixic Acid (CNA) agar plates under 5% CO_2_.

To assess the impact of residual nutrients from rich media on bacterial growth in nutrient-limited conditions, strains of *E. coli* (UMB3190, aka ATCC 10798) and *Klebsiella pneumoniae* (UMB9987) were first grown overnight in BHI. These precultures then were diluted 1:100 into either mMP-AU or M9 salts and cultured under our standard conditions: 48 hours at 5% CO_2_, 37°C with orbital agitation of 200 rpm.

### SimUrine Formulation Optimization

Several versions of SimUrine were tested and the final formulation is described in **Supplementary Protocols 1** and **Supplementary Table 2**. Other versions are included for reference in **Supplementary Protocols 2.** These formulations were designed to mimic the nutrient-limited bladder environment through progressive enrichment of mMP-AU and by tuning reagent concentrations to stay within urine ranges, while designing a modular protocol to enable facility, prolonged storage, and component replacement. Different formulations were tested for growth of selected isolates, previously cultured on BHI agar plates.

The first version of SimUrine (**SimUrine.v1**) was formulated as follows: mMP-AU supplemented by adding N-acetylglucosamine, L-threonine, and L-serine (to mimic mucin composition without the turbidity or contamination problems associated with direct use of mucins) and lactic acid, pyruvic acid, acetic acid, manganese sulfate, and L-cysteine (to protect facultative species and improve fastidious species growth). Bacterial growth was assessed for two consecutive periods of 48 hours.

Based on the performance of **SimUrine.v1**, and with the aim to extend use of the medium to a wider range of microbes, a second version of SimUrine was designed (**SimUrine.v2**). Inspired by the work on bacterial communities from the gut (Shetty et al., 2022), we incorporated vitamins and trace elements into this formulation.

Given the biological relevance of iron, we assessed the impact of hemin supplementation in our formulation with two concentrations: one as described in **SimUrine.v3** and one tenfold greater. The need of uric acid for bacterial growth also was evaluated by growing strains in the presence and absence of this compound.

To reduce spontaneous precipitate formation, improve medium stability, and keep uric acid in our formulation, the reagent order was modified in **SimUrine.v4**: now uric acid was added after urea and not before. We also improved buffering capacity of the medium with the addition of 4-(2-hydroxyethyl)-1-piperazineethanesulfonic acid (HEPES), and to better mimic the nutrient profile of urine, we added non-ionic surfactant Tween80 and extra peptides (Reitzer & Zimmern, 2019; Schmied et al., 2021). Given the role of cysteine as a key energy and carbon source in various environments (Göbbels et al., 2021), we also assessed the use of its dimeric form cystine, by testing a version of **SimUrine.v.4** with cysteine replaced by cystine.

To minimize precipitate formation during storage, and reduce chelation of trace elements, sodium bicarbonate and 3-(N-morpholino) propanesulfonic acid (MOPS) were added as buffering agents instead of HEPES, resulting in **SimUrine.v5**. In contrast to earlier formulations that included autoclaving of the final medium, this new formulation was sterilized by filtration with a 0.2 µm filter.

To optimize conditions for *Lactobacillus* species growth, while considering the biological relevance of the following nutrients, **SimUrine.v5** was enriched with glucose, FeSO_4_, L-valine, L-threonine, and L-tryptophan (Kwoji et al., 2022; Liu et al., 2020; Ramoneda et al., 2023; Saguir & De Nadra, 2007; Veith et al., 2015). NaHCO_3_ was added and pH reduced to increase stability. Three different concentrations of urea were evaluated. Based on these experiments, a final version, **SimUrine.v6** version was adopted.

### SimUrine Evaluation

To assess the performance of the different SimUrine versions, pure bacterial isolates were first grown overnight in a 20 ml glass tube containing 4 ml of the corresponding version of SimUrine at standard conditions (37°C, 5% CO_2_ with orbital agitation at 200 rpm). Overnight cultures were diluted 1:100 into the noted SimUrine versions, and growth was monitored in 96-well flat-bottom plates under standard conditions (37°C, 5% CO_2_ and orbital agitation at 200 rpm) for 24, 48 or 72 hours (depending on the species). Optical density (OD) was assessed every ten minutes using Biotek (Winooski, VT, USA) or Cerillo Alto (Charlottesville, VA, USA) plate readers (indicated with each result). To confirm species identity, colonies were obtained from the cultures at the end of each growth curve and identified using mass spectrometry on a Bruker MALDI-TOF (Billerica, MA, USA).

### Physicochemical Properties of SimUrine.v6

To characterize **SimUrine.v6** for broader applications, physicochemical parameters (density, refractive index, viscosity, conductivity, and osmolarity) were determined from two independent batches prepared two weeks apart, each measured in technical triplicate at 20°C. Density was determined using a volumetric flask and an analytical balance, refractive index was measured using a KERN Abbe ORT 1RS refractometer (Kern & Sohn, Balingen, Germany), viscosity using an MCR 702e rheometer (Anton Paar, Breda, The Netherlands), conductivity using a portable conductivity meter (Mettler-Toledo, Columbus, OH, USA) and osmolarity using an Omo1 osmometer (Advance instruments, Norwood, MA, USA).

### Evaluation of SimUrine.v6 in Microbial Ecology-Relevant Experiments

To compare the performance of **SimUrine.v6** for urobiome-related assays, antibiotic susceptibility tests were performed on *E. coli* (UPEC20). A 1:19 ratio of Trimethoprim (TMP) and sulfamethoxazole (SMX) was tested at a starting concentration of 32 µg/mL TMP and 608 µg/mL SMX, with serial two-fold dilutions. Susceptibility to kanamycin was also tested, starting at 128 µg/mL. MHB was used as the reference medium. In all assays, the initial inoculum was ∼ 2.5 x 10^5^ CFU/mL. Optical density was measured in a 96-well plate using a Cerillo Alto reader (Cerillo, Charlottesville, VA) at 37°C and 5% CO_2_.

### Co-culture of *E. coli* and *E. faecalis* Strains

Reference strains of *E. coli* (ATCC25922) and *E. faecalis* (ATCC29212) were obtained from the American Type Culture Collection (ATCC). Prior to experimental use, both strains were sub-cultured onto Trypticase Soy Agar (TSA) supplemented with 5% sheep blood and incubated at 37°C in a 5% CO₂ atmosphere for bacterial propagation and viability assessment. Overnight cultures of both bacterial strains were prepared in NYCIII medium and SimUrine.v6 medium, with a minimum of five biological replicates per strain. Cultures were incubated at 37°C with orbital shaking at 200 rpm. Following overnight incubation, cultures were harvested by centrifugation at 3,000 × g for 10 minutes at room temperature, supernatants were discarded, and bacterial pellets were resuspended in fresh medium (NYCIII or SimUrine.v6, respectively). Bacterial cell density was standardized using optical density measurements at 600 nm (OD₆₀₀), with all cultures adjusted to an initial OD₆₀₀ of 0.05. For growth kinetic measurements, 80 μL of standardized bacterial suspension was combined with 720 μL of the appropriate growth medium (NYCIII or SimUrine.v6) in individual wells of a dual-well system (Cerillo co-culture duet system, Charlottesville, VA). Control wells contained 800 μL of sterile medium without bacterial inoculum.

Growth kinetics were monitored over 24 hours using a Cerillo Alto microplate reader with continuous orbital shaking at 200 rpm at 37°C. Optical density measurements (OD₆₀₀) were recorded at 30-minute intervals, and final readings at 24 hours were used for subsequent data analysis. Growth curves and standard error calculations were performed using MATLAB-R2025a (The MathWorks, Inc., 2025).

### SimUrine.v6 Protocol for 500 mL of Media (Abbreviated)

#### Preparation

Follow this abbreviated protocol in a sterile environment using aseptic technique to ensure reproducibility. All stock solutions must be prepared and either filtered (0.2 µm) or autoclaved separately before making the medium (see **Supplementary Protocol 1 and Supplementary Table 2, for details and reagents**).

Once stock solutions are prepared and sterilized, dissolve the module reagents sequentially as indicated into 200 mL of dH_2_O, stirring at 40°C: starting with the **mMP-AU** module, followed by the **Amino acids** module, the **Carbon sources** module, and the **Others** module. While the medium is warm, add 500 µL of Tween80, 1.05 g of MOPS and 7.5 g of urea. Continue adding the stock solutions sequentially, check and adjust pH to 6 with HCl or NaOH, add freshly prepared uric acid solution, and finally L-serine, as indicated in the protocol. Check pH and adjust if needed. Complete volume to 500 mL with H_2_O. Filter-sterilize the media using a 0.2 µm filter. To avoid precipitation, avoid storage at temperatures lower than 4°C. To extend lifespan, protect from light.

## RESULTS

### Optimization of mMP-AU for Bacterial Survival

To compare viability of *E. coli* (UMB1180) in minimal salt bases, we monitored survival over 8 days. Survival was marginally better in mMP-AU compared to M9 salts across the whole experiment (**Fig.1**). This trend was maintained through day 8, indicating that mMP-AU supports bacterial viability at least as effectively as M9.

**Fig. 1.**
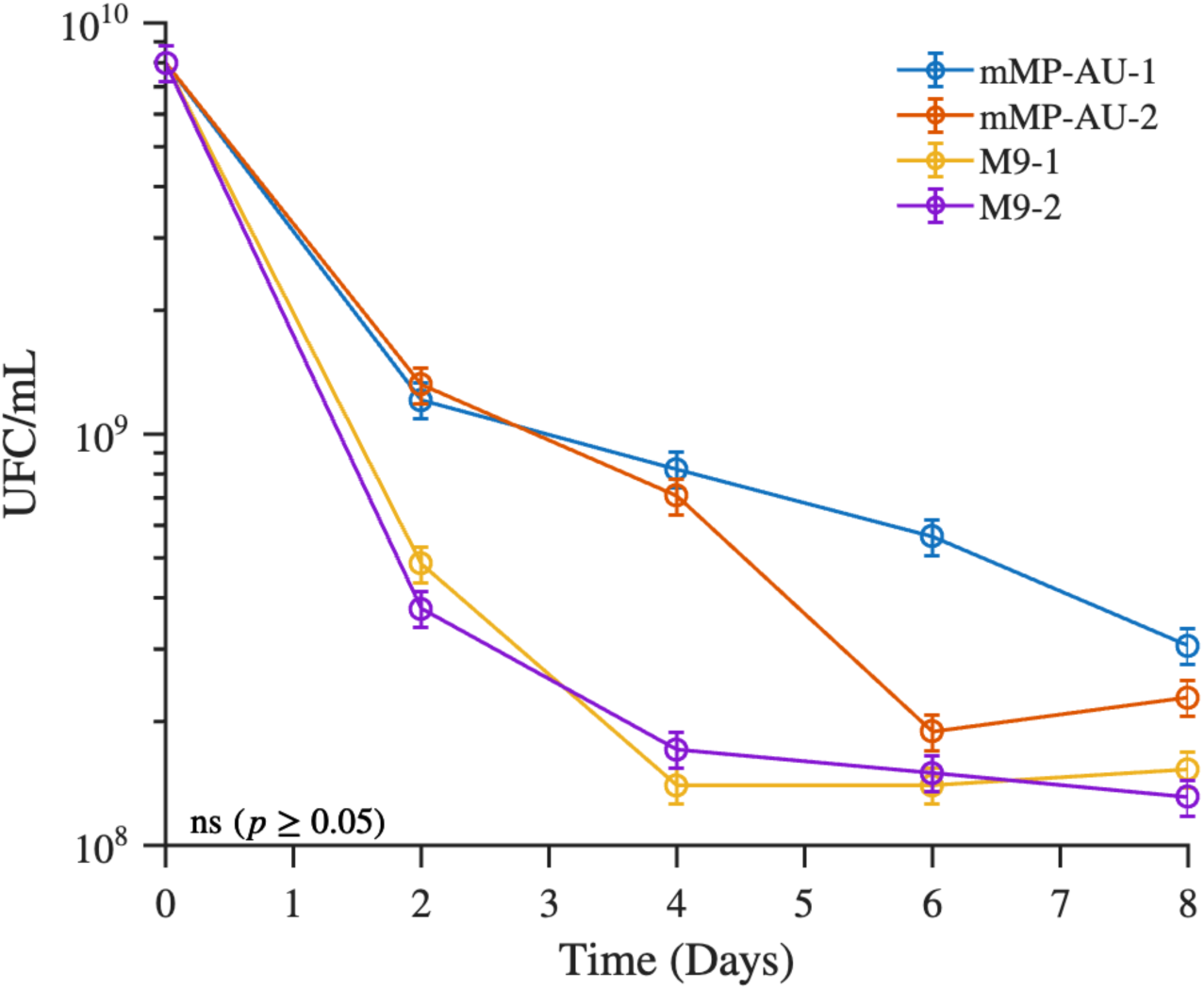
Survival of *E. coli* (UMB1180) in either mMP-AU or M9 salts. Two technical replicates were conducted for each condition, with each replicate averaging two biological replicates. The y-axis displays logarithmic value of CFU/mL of *E. coli,* and the x-axis represents time in days, standard deviation is displayed for each point. These results suggest that both salt bases similarly support limited survival of urobiome species.

### Glucose Supplementation Influences Survival of Urinary Isolates

To evaluate the contribution of a carbon source to survival of urobiome species, we incorporated glucose to the minimal salt bases. For mMP-AU and M9 salts, glucose supplementation had a contrasting effect on survival of a urinary isolate of *Lactobacillus jensenii* (UBM8651). For mMP-AU, a glucose concentration above 2% reduced survival, whereas elevated glucose concentrations had little effect on survival in M9 salts (**Supplementary Figure 1A**); however, survival on either salt base did not extend beyond one day. For *Aerococcus urinae* (UBM5254), increased glucose concentration was directly associated with enhanced survival when using mMP-AU as the salt base. In contrast to *L. jensenii* (UBM8651), *A. urinae* (UBM5254) survived for a second day, but only in glucose-supplemented M9 salts (**Supplementary Figure 1B**).

As elevated glucose concentrations reduced *L. jensenii* (UMB8651) survival in mMP-AU, we tested the impact of various carbon sources and nutrients on UMB8651 growth in modified versions of mMP-AU. Growth was primarily supported by amino acids and vitamins, while other supplements failed to fully sustain bacterial proliferation (**Supplementary Figure 2**).

### Effect of Residual Nutrients from the Inoculum on Bacterial Growth

Strains of *E. coli* (UBM3190) and *K. pneumoniae* (UBM9987) pre-grown on the rich medium BHI and then transferred to mMP-AU or M9 salts without supplementation reached between one-tenth and one-third of their maximum OD when grown in BHI, indicating that residual nutrients from the rich medium supported partial growth (**Fig.2A and Fig.2B**).

**Fig. 2.**
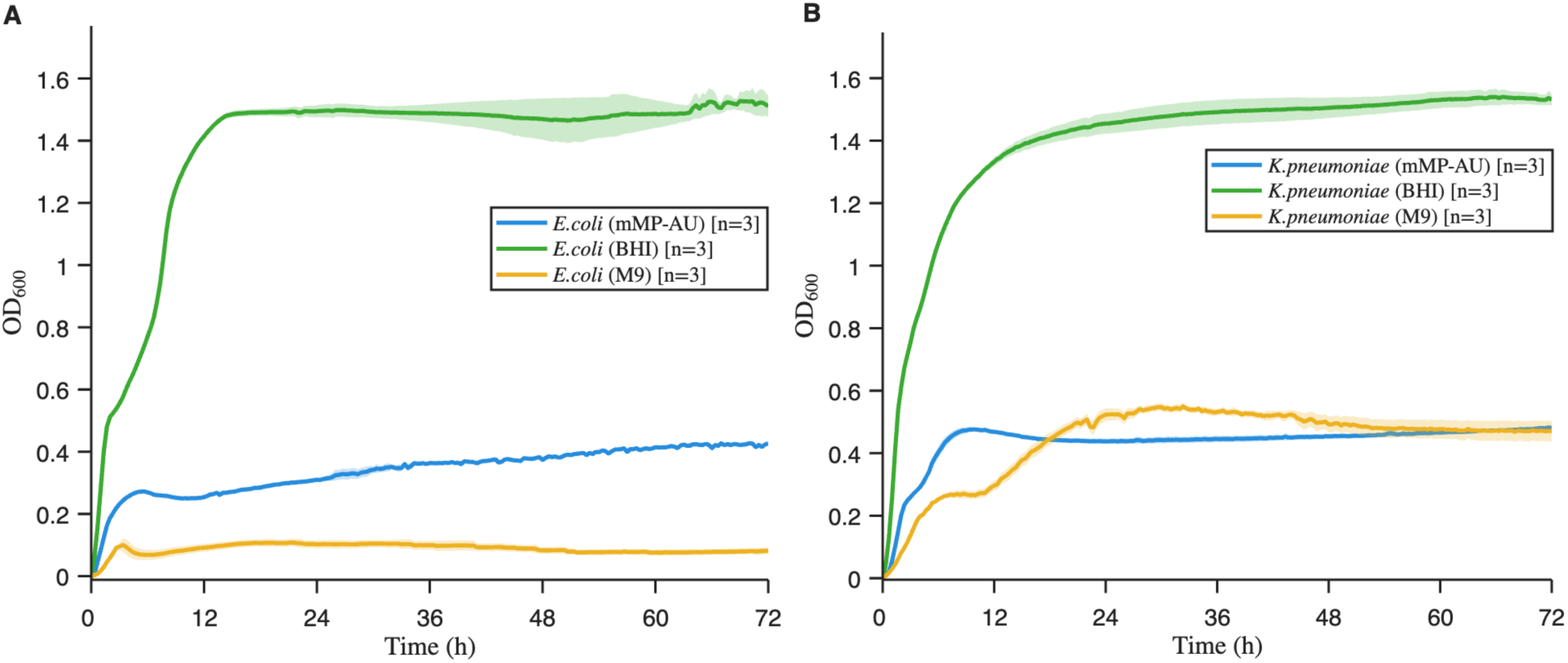
Effect of trace nutrients from complex media on bacterial growth. Overnight cultures of *E. coli* (UMB3190) (A) and *Klebsiella pneumoniae* (UMB9987) (B) were diluted 1:100 into BHI, M9 or mMP-AU and incubated for 48 hours. Optical density at 600 nm (OD_600_) is shown on the y-axes; time (hours) on the x-axes. Solid lines represent mean values and shading standard error of the mean.

### SimUrine Formulation Optimization

#### SimUrine.v1

This simplest version of SimUrine supported the discrete growth of the facultative anaerobic bacterial species *E. coli* (UBM3190), *Actinotignum schaalii* (UBM13319), *A. urinae* (UBM5254), *Klebsiella pneumoniae* (UBM9987) and *Streptococcus anginosus* (UBM8616), while failing to support the growth of *Enterococcus faecalis* (**Fig.3**).

**Fig. 3.**
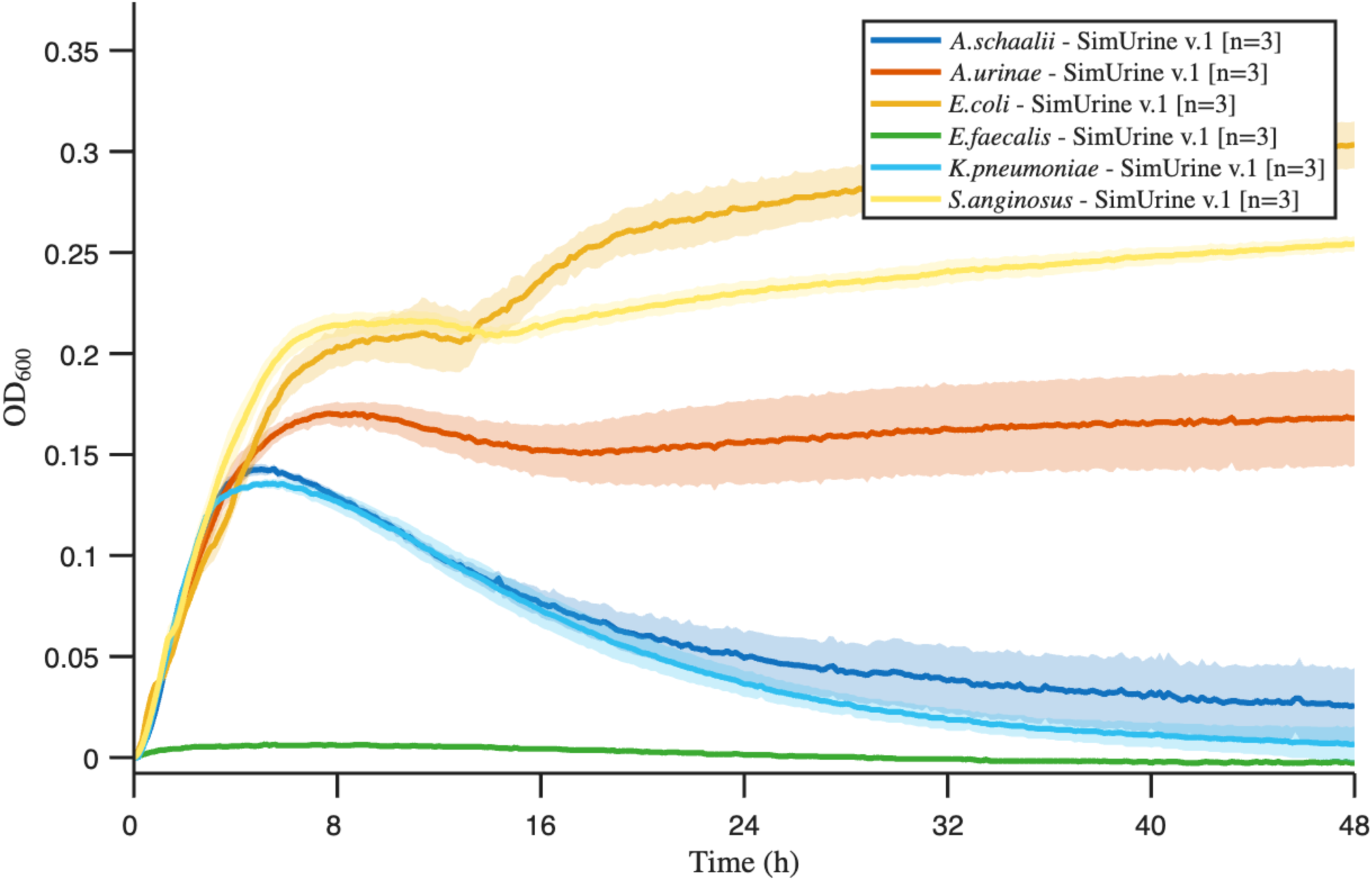
Growth in SimUrine.v1. Cultures of different bacterial isolates over 48 hours. OD_600_ is shown on the y-axis; time (hours) on the x-axis. Solid lines represent mean values and shading standard error of the mean.

#### SimUrine.v2

Adding vitamins and trace elements to our formulation markedly enhanced the growth of *E. coli* (UBM3190), with OD_600_ values increasing from 0.2-0.3 (**Fig.3**) to 0.7, which exceeded growth in BHI (**Fig.4A**). *Klebsiella pneumoniae* (UBM9987) also showed improved growth compared to SimUrine.v.1 (**Fig.3 and Fig.4B**); however, unlike *E. coli* (UBM3190), *K. pneumoniae* did not surpass its growth in BHI. Despite enhanced bacterial growth with this version, a precipitate formed in SimUrine.v2 after 24 hours of preparation (**Supplementary Figure 3**).

**Fig. 4.**
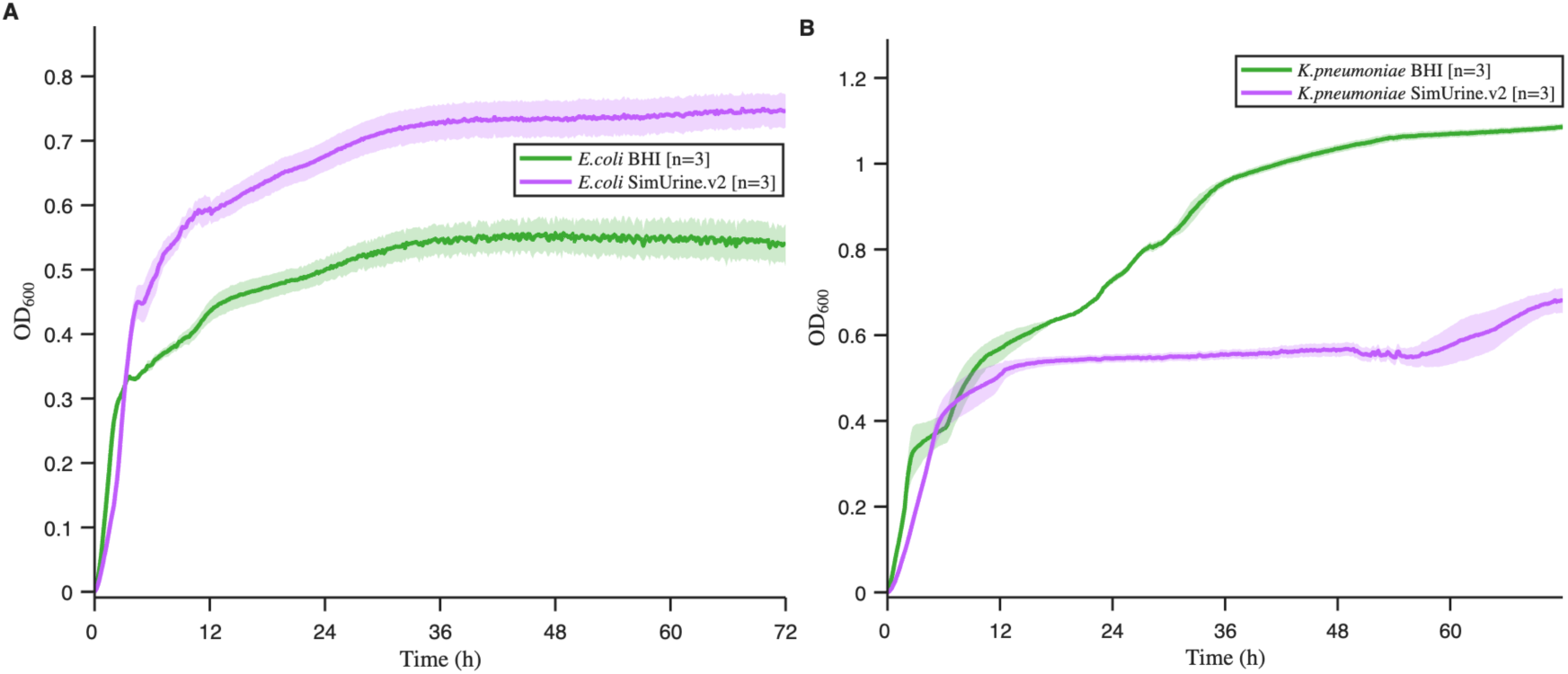
Growth in SimUrine.v2. Cultures of *E. coli* (UBM3190) (A) and *K. pneumoniae* (UMB9987) (B) in SimUrine.v2. Optical density at 600 nm (OD_600_) is shown on the y-axis; time (hours) on the x-axis. Parallel cultures in BHI (green lines) were used as a reference. Solid lines represent mean values and shading standard error.

#### SimUrine.v3

Hemin supplementation (either 1X or 10X) to produce SimUrine.v3 did not substantially improve *E. coli* CFT073 growth (compared to SimUrine.v2 growth) but had a marked effect on the growth characteristics of *S. anginosus* (UBM8616) (**Fig.5A and Fig.5B**). Based on these results, we adapted our formulation, reducing the concentration of hemin from 10X to 5X, while evaluating the importance of uric acid for bacterial growth. These modifications revealed that uric acid was essential for reaching higher OD_600_ values and enhancing exponential growth, while hemin 5X performed well (**Supplementary Figure 4**). Hemin incorporation defined SimUrine.v.3.

**Fig. 5.**
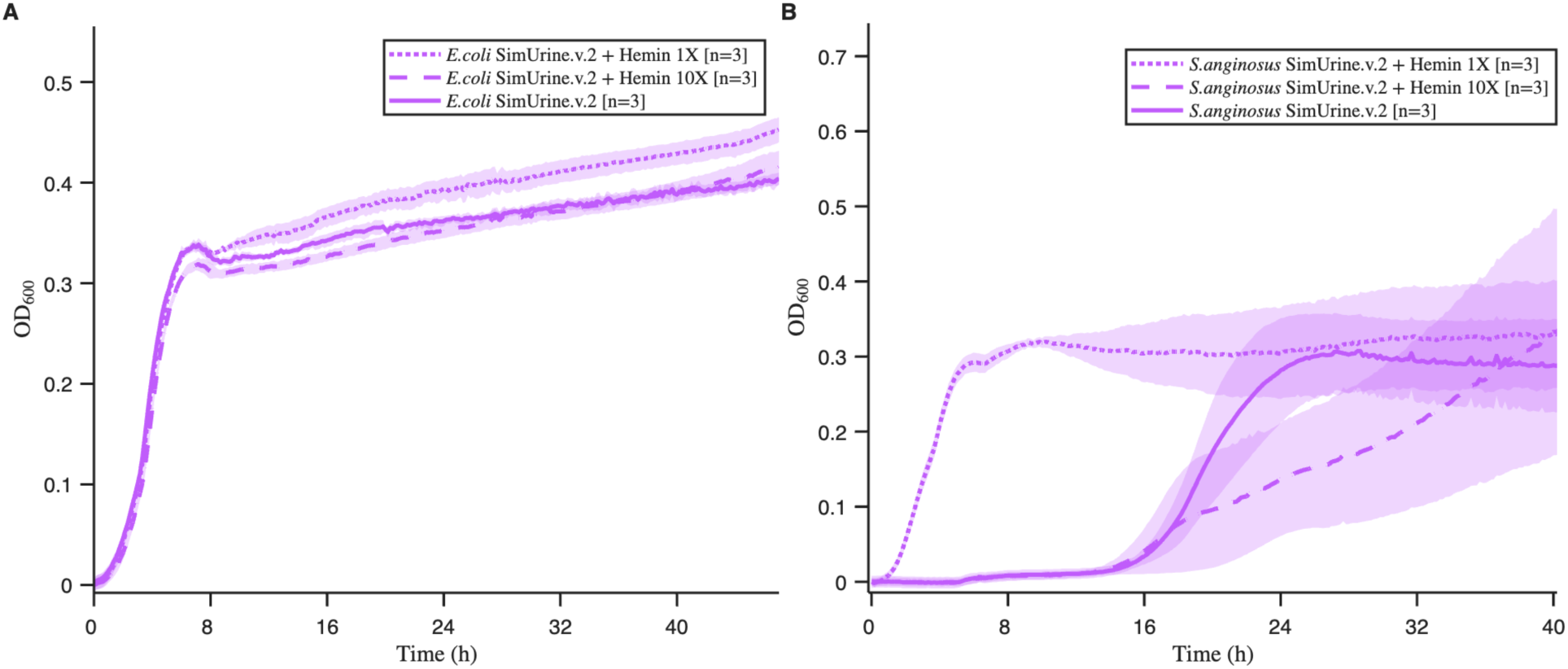
Growth in SimUrine.v2 and hemin supplemented versions. Cultures of *E. coli* (UBM3190) (A) and *S. anginosus* (UMB8616) (B) in SimUrine.v2 (solid lines) and SimUrine.v.2 supplemented with hemin 1X (dotted lines) and hemin 10X (dashed lines). Optical density at 600 nm (OD_600_) is shown on the y-axes; time (hours) on the x-axes. Lines represent mean values and shading standard error of the mean.

#### SimUrine.v4

Represented a major improvement of SimUrine formulation. We incorporated amino acids within urine-relevant concentration ranges (Schmied et al., 2021), changed the order of hemin and uric acid supplementation, and introduced both HEPES and Tween80. We used this formulation to test if bacteria could grow with cysteine or its dimer, cystine. For *E. coli* (UBM3190) and *K. pneumoniae* (UBM9987), SimUrine.v4 resulted in vastly improved OD_600_ (compare **Fig.6A and Fig.6B** to **Fig.5A and Fig.5B)** and resulted in substantial growth of *E. faecalis* and *C. riegelii* (**Fig.6C and Fig.6D**). In all cases, cysteine performed better than cystine.

**Fig. 6.**
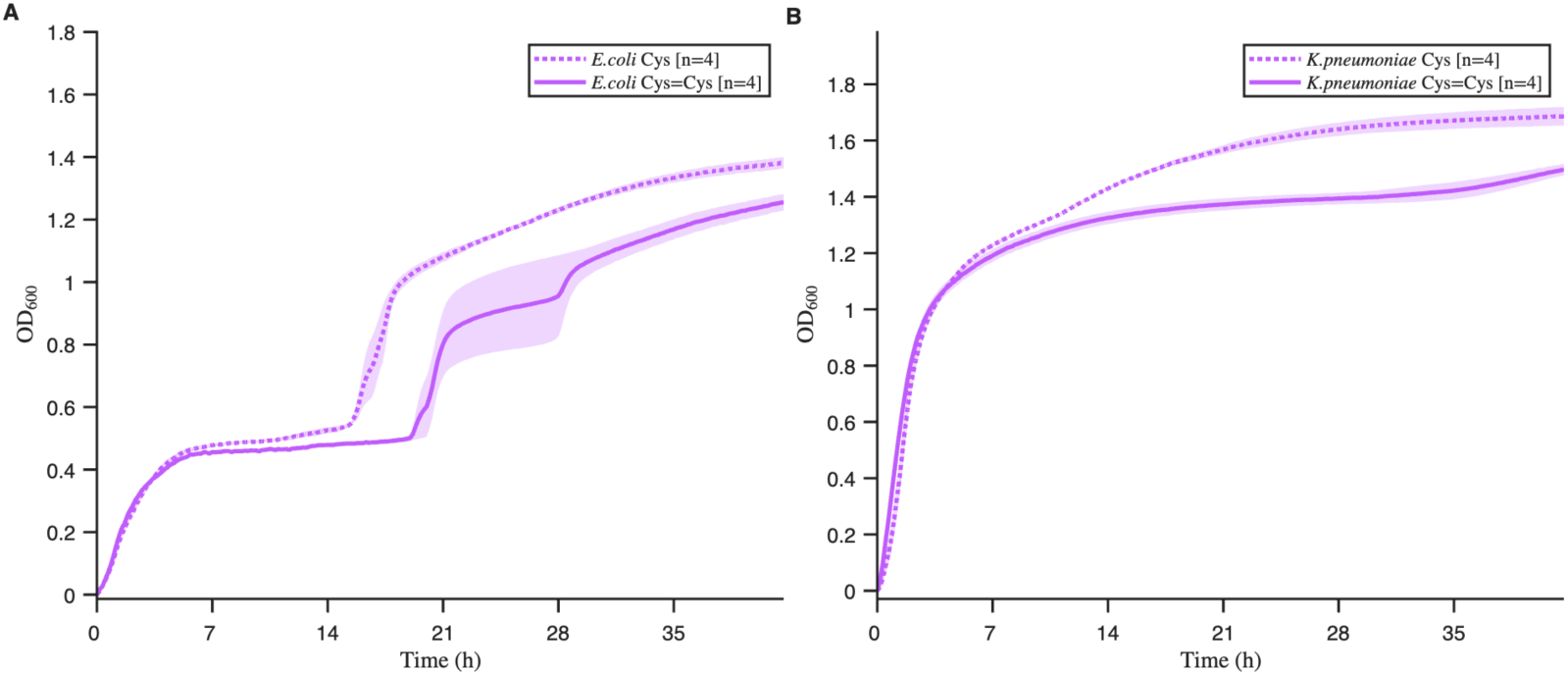
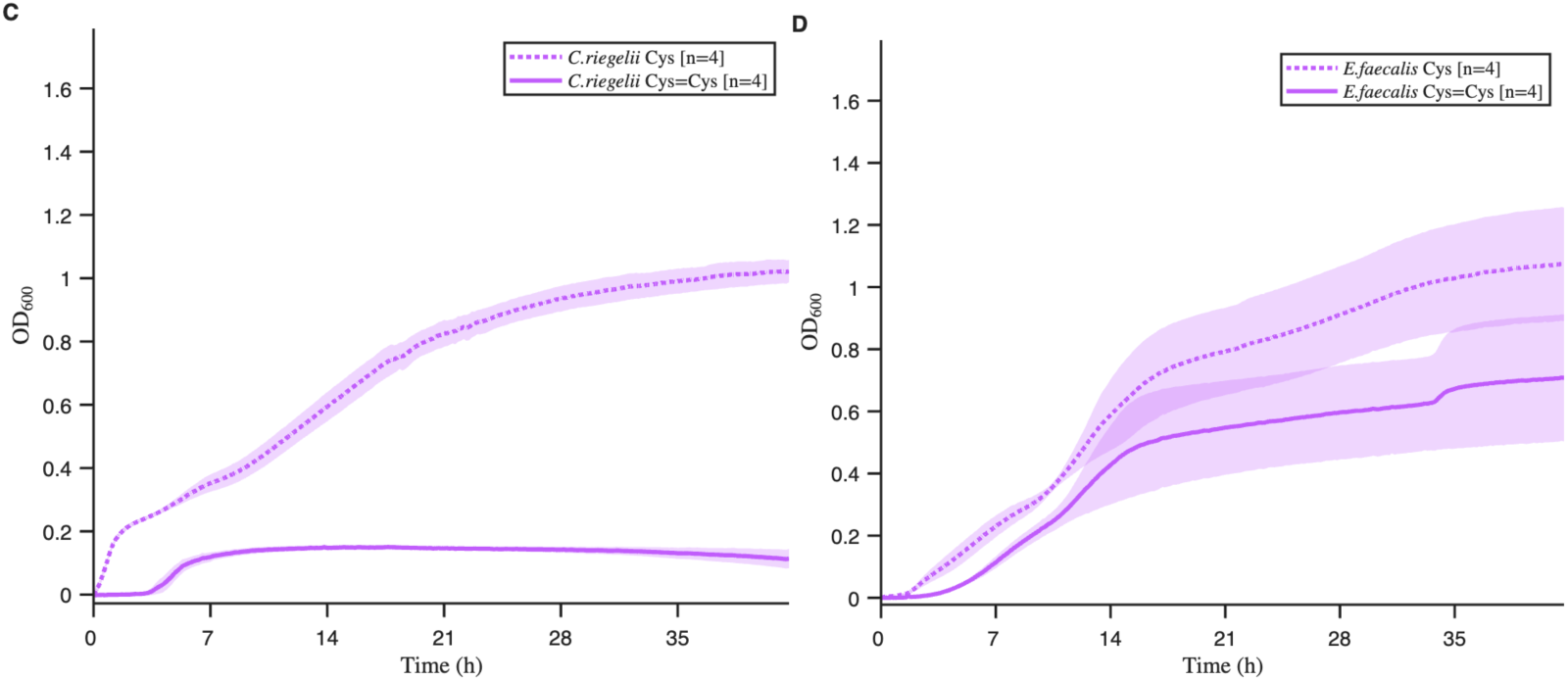
Growth in SimUrine.v4. Cultures of *E. coli* (UBM3190) (A), *K. pneumoniae* (UBM9987) (B), *C. riegelii* (UBM12267) (C) and *E. faecalis* (UBM3193) (D) in cysteine (dotted lines) or cystine (solid lines). Optical density at 600 nm (OD_600_) is shown on the y-axes; time (hours) on the x-axes. Lines represent mean values and shading standard error of the mean.

#### SimUrine.v5

We increased long term formulation stability without altering growth characteristics by replacing HEPES with MOPS, incorporating sodium bicarbonate, and by filter-sterilizing the medium (data not shown).

#### SimUrine.v6

Supplementation of SimUrine.v5 with glucose, FeSO_4_, L-valine, L-threonine, and L-tryptophan supported the growth of a larger number of microbes than previous versions, including *Lactobacillus* species, *E. faecalis, Proteus mirabilis, Prevotella bucallis,* among others (**Supplementary Table 1**). Urea concentration did not substantially affect the growth of *E. faecalis* (UBM3193) (**Fig.7A**), but the higher urea concentrations improved the growth of *P. mirabilis* (UBM7310) (**Fig.7B**). The new formulation reduced growth of *E. faecalis* compared to SimUrine.v4 (**Fig.6D**). Despite the reduced growth, the higher urea concentration was kept in the final formulation to better reflect physiological urine levels, as bacterial viability was not compromised.

**Fig. 7.**
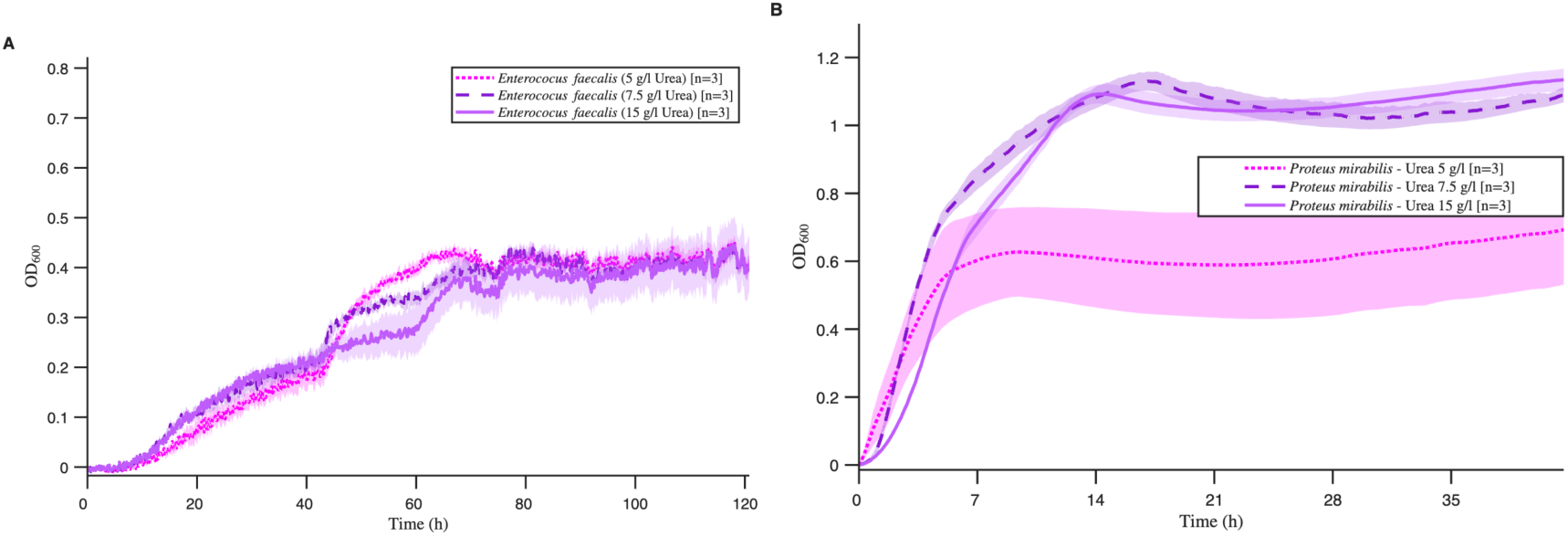
Effect of urea in SimUrineV6. *E. faecalis* (UBM7540) (A), and *P. mirabilis* (strain 83) (B) in SimUrineV6 with 3 different concentrations of urea. Optical density at 600 nm (OD_600_) is shown on the y-axes; time (hours) on the x-axes. Lines represent mean values and shading standard error.

### Use of SimUrine.v6 in biologically relevant urobiome microbial experiments

We evaluated the use of SimUrine.v6 in two ecology-relevant contexts: antibiotic susceptibility testing and a bacterial interaction assay. *E. coli* (UPEC20) exhibited reduced susceptibility to both TMP-SMX and kanamycin in SimUrine.v6 (**Fig.8A and Fig.9A**), compared to that observed in the richer, complex reference medium, MHB (**Fig.8B and Fig.9B**).

**Fig. 8.**
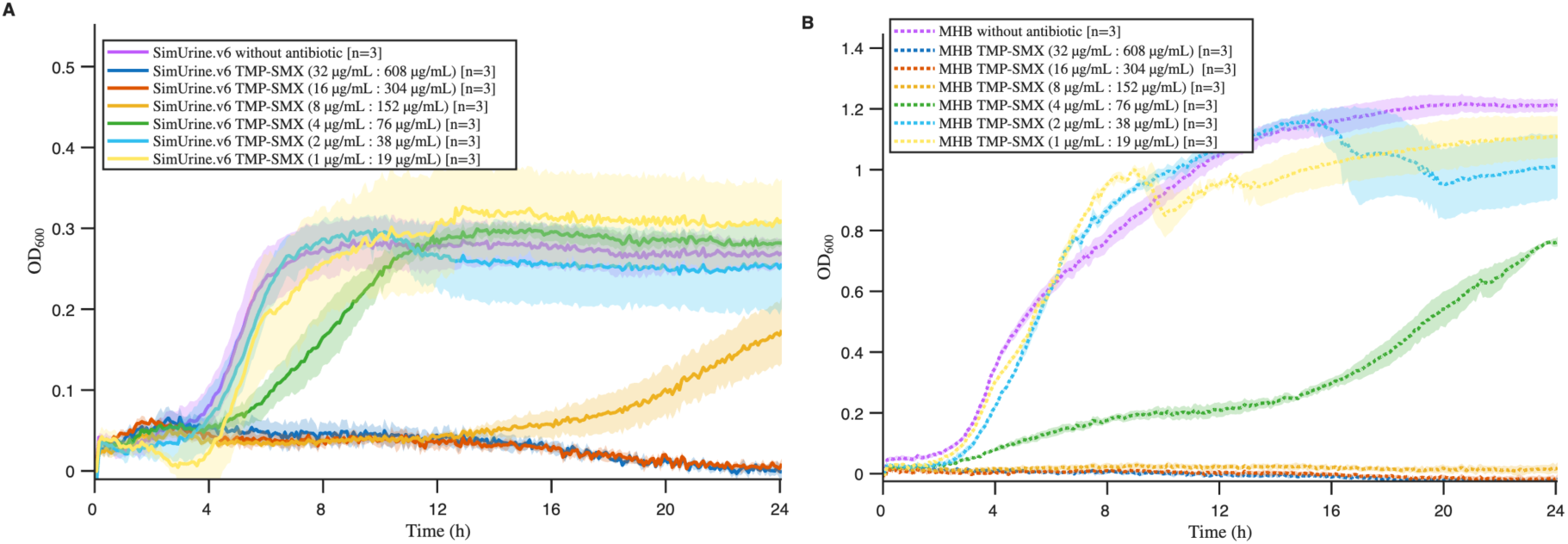
TMP-SMX susceptibility of *E. coli* (UPEC20). Cultures of *E. coli* in SimUrine.v6 (A) and MHB (B). Initial antibiotic concentration TMP-SMX (Trimethoprim 32 µg/mL: Sulfamethoxazole 608 µg/mL) was serially diluted by half each time, to a final concentration of TMP-SMX 1µg/mL:19 µg/mL. Cultures in media without antibiotic were used as control. Optical density at 600 nm (OD_600_) is shown on the y-axis; time (hours) on the x-axis. Lines represent mean values and shading standard error of the mean.

**Figure 9.**
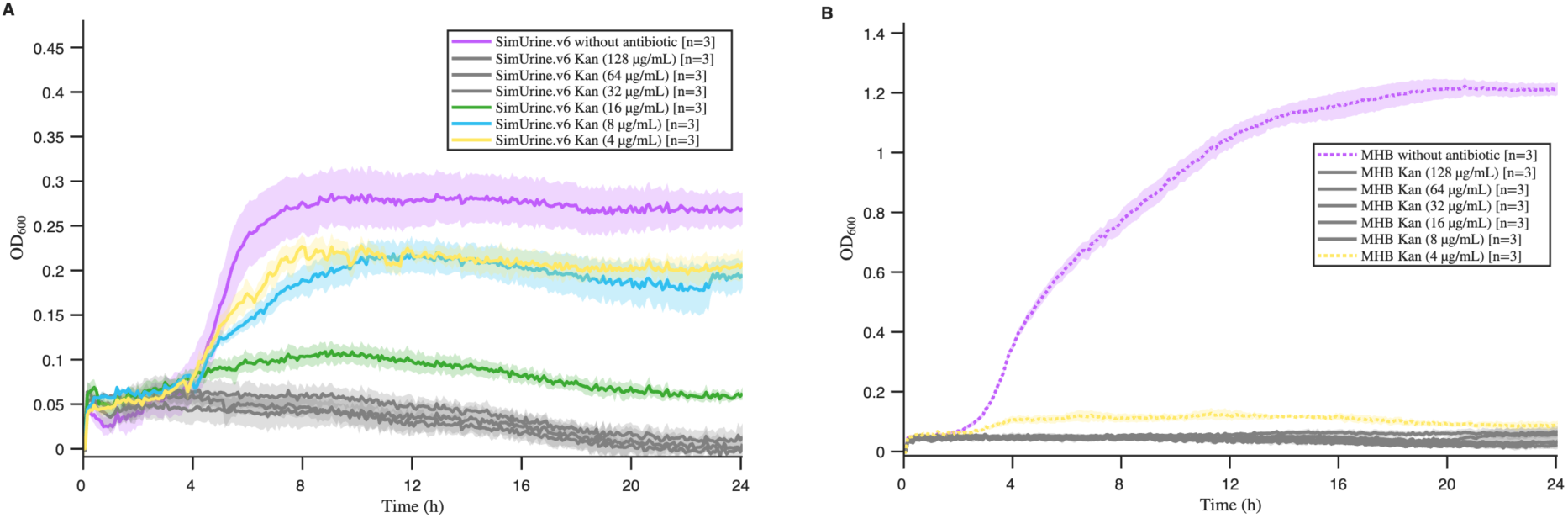
Kanamycin susceptibility of *E. coli* (UPEC20). Cultures of *E. coli* in SimUrine.v6 (A) and MHB (B) in decreasing concentrations of Kanamycin. Initial antibiotic concentration Kan (128 µg/mL) was serially diluted by half each time, to a final concentration of Kan (4µg/mL). Gray curves indicate no growth for both media; colored curves indicate growth. Cultures in media without antibiotic were used as control. Optical density at 600 nm (OD_600_) is shown on the y-axis; time (hours) on the x-axis. Lines represent mean values and shading standard error of the mean.

Growth characteristics of *E. coli* (ATCC 25922) and *E. faecalis* (ATCC 29212) also differed in SimUrine.v6 versus the rich, complex NYCIII. In NYCIII, entry of *E. coli* into exponential growth was rapid and there was little difference in the growth characteristics of *E. coli* cultured alone and in co-culture with *E. faecalis* (**Fig.10A**). In SimUrine.v6, entry of *E. coli* into exponential growth was similarly slower when grown either alone or in co-culture with *E. faecalis*, but entry of *E. coli* into stationary phase was more abrupt and occurred at a lower OD when co-cultured with *E. faecalis* (**Fig.11A**). In NYCIII, entry of *E. faecalis* into exponential growth was similarly rapid when grown alone or in co-culture with *E. coli*, while entry into stationary phase occurred at a slightly lower OD when co-cultured with *E. coli* than when grown alone (**Fig.10B**). In SimUrine.v6, *E. faecalis* grown alone experienced a long lag but when co-cultured with *E. coli* failed to grow (**Fig.11B**).

### SimUrine.v6 physicochemical parameters

SimUrine.v6 exhibited physicochemical properties within the normal range of human urine (as described in the literature (**Table 1**), including low viscosity, making it suitable for microfluidic applications.

**Table 1.**
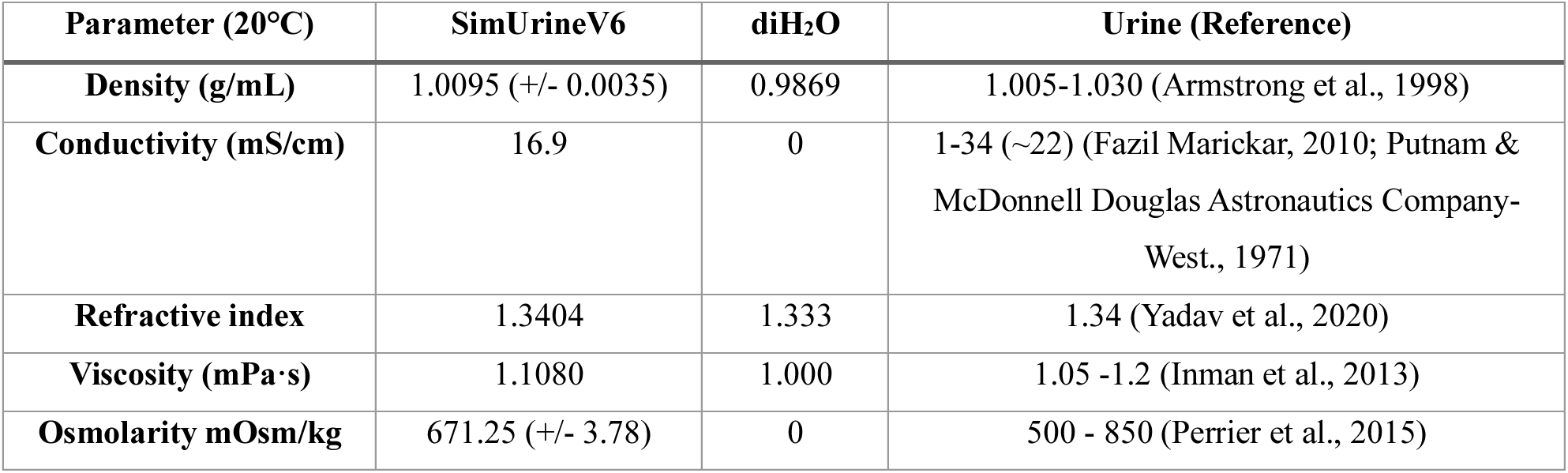

## DISCUSSION

Conventional culture media for human urobiome studies utilize standardized formulations that fail to reflect the urinary tract’s natural biochemical environment. This discrepancy stems primarily from the use of undefined reagents that create complex, proprietary formulations that are difficult to replicate across laboratories. The human bladder maintains distinct physiological parameters including specific pH, osmolarity, and metabolite concentrations that differ substantially from commercial media compositions. We developed SimUrine, a fully chemically defined medium where each component was selected to either match the documented human biochemical parameters or support urobiome growth at minimal effective concentrations.

Following urobiome discovery, researchers have attempted to isolate, culture and characterize these microbes, using artificial urine media or enriched formulations (e.g., NYCIII medium) to maximize isolate recovery. While these approaches have enabled the isolation and study of novel strains, current media formulations have limited our understanding of their behavior under native conditions.

SimUrine development integrates current knowledge of urine composition, artificial urine media, and cultivation strategies used for other microbial communities, such as the gut microbiome (Shetty et al., 2022). By adapting the established MP-AU medium, we successfully cultured bacteria important for bladder health and disease while providing an open, defined, and modifiable medium formulation. This medium permits the behavioral study of many fastidious and anaerobic bacteria species that previously failed to survive under previous strategies.

Development of SimUrine followed a systematic approach, beginning with a basic salt base (MP-AU) that was subsequently enriched with multiple carbon sources to identify the most relevant biochemical requirements for bacterial growth. Our initial version (SimUrine.v1) incorporated nutrients accessible to the urobiome within the bladder environment. While N-acetylglucosamine, L-threonine, and L-serine may not be components of excreted urine, the urobiome has access to mucins and epithelial cells within the bladder lining, from where these nutrients can be metabolized (Neugent et al., 2023). This formulation yielded improved growth for the tested bacteria (**Fig.3**).

To enable growth of a broader range of microbes, we selected nutrients and vitamins identified from successful gut microbiome cultivation studies. SimUrine.v2 included common vitamins and trace elements, resulting in enhanced growth of *E. coli* (UBM3190) and *K. pneumoniae* (UBM9987) (**Fig.4**), although it did not support the optimal growth of other species of interest, such as *E. faecalis* and *A. urinae*, among others. Subsequent testing revealed that hemin incorporation and expanded amino acid availability had considerable effects on bacterial growth, as demonstrated in SimUrine.v.4 (**Fig.5, Supplementary Figure 4**).

The inclusion of Tween80, an important medium surfactant, along with improved buffering capacity through HEPES addition, extended media utility to more fastidious organisms such as *C. riegelii* (**Fig.6**). However, *Lactobacillus* species remained uncultured with this formulation. Final optimization involved replacing HEPES with MOPS, adding sodium bicarbonate, implementing filter sterilization rather than autoclaving, and supplementing with FeSO_4_, which enabled *Lactobacillus* to grow in glass tubes (**Supplementary Table 1**). We hypothesize that HEPES combined with phosphate buffer provides insufficient buffering capacity under 5% CO_2_ incubation, potentially contributing to trace element chelation that affects fastidious bacterial growth. In contrast, MOPS maintains pH stability and reduces trace element chelation, while FeSO_4_ enhances iron bioavailability. Whereas we could culture *Lactobacillus* species in glass tubes, we have not yet been able to obtain reliable growth curves using 96-well plates. Increasing the diameter and volume of the wells may enable reliable growth curve generation.

Beyond the logical importance of using a medium that better replicates actual urine composition, our antibiotic susceptibility and co-culture results demonstrate the feasibility of utilizing SimUrine in clinical investigations. Using this fully defined medium, for the tested strain (**Fig.8**), we detected lower antibiotic susceptibility than standard values for *E.coli* (Lewis II et al., 2023), even when using *in vivo*-like TMP-SMX proportions (U.S. Food and Drug Administration, 2011), potentially reflecting susceptibilities closer to those encountered under native bladder conditions. Additionally, SimUrine enabled observation of previously unknown interactions between urobiome community members, highlighting the importance of physiologically relevant culture conditions for understanding clinical microbiology.

SimUrine enables the use of physiological urea concentrations without completely suppressing growth of urea-susceptible bacteria (**Fig.7**). Urea incorporation proved beneficial for species such as *P. mirabilis* (**Fig.7B**). Indeed, urea is noted to be the most abundant organic molecule in human urine at concentration ranges between 9.3-23.3g/L (Putnam & McDonnell Douglas Astronautics Company-West, 1971). At 15g/L, SimUrine’s urea concentration allows for more accurate representation of a natural bladder environment while maintaining broad bacterial compatibility.

SimUrine represents a fully defined culture medium that supports growth of diverse bacterial species and clinical isolates while remaining adaptable to specific research and clinical requirements. The medium’s complete chemical definition ensures reproducibility across laboratories, while its modular design allows targeted modifications without compromising stability. Previous artificial urine compositions, such as those formulated by Brooks and Keevil or Zandbergen, utilize protein hydrolysates that provide wide coverage of necessary amino acids, but introduce unnecessary variability via undefined compositions (Brooks & Keevil, 1997; Zandbergen et al., 2021). With defined nutrient compositions, the dietary behaviors of tested bacteria can be fully characterized with SimUrine.

The clinical implications of this work extend beyond improved pathogen recovery and characterization. By providing culture conditions that more accurately reflect the native bladder environment, SimUrine enables more clinically relevant assessment of antimicrobial susceptibility patterns (**Fig. 8 and Fig.9**) and microbial interactions (**Fig.11**) that may be not observed when using rich media (**Fig.10**) (Lewis II et al., 2023). We acknowledge that the *E. faecalis* suppression in co-culture (**Fig.11B**) needs mechanistic investigation, which exceeds the purpose of this work.

**Fig. 10:**
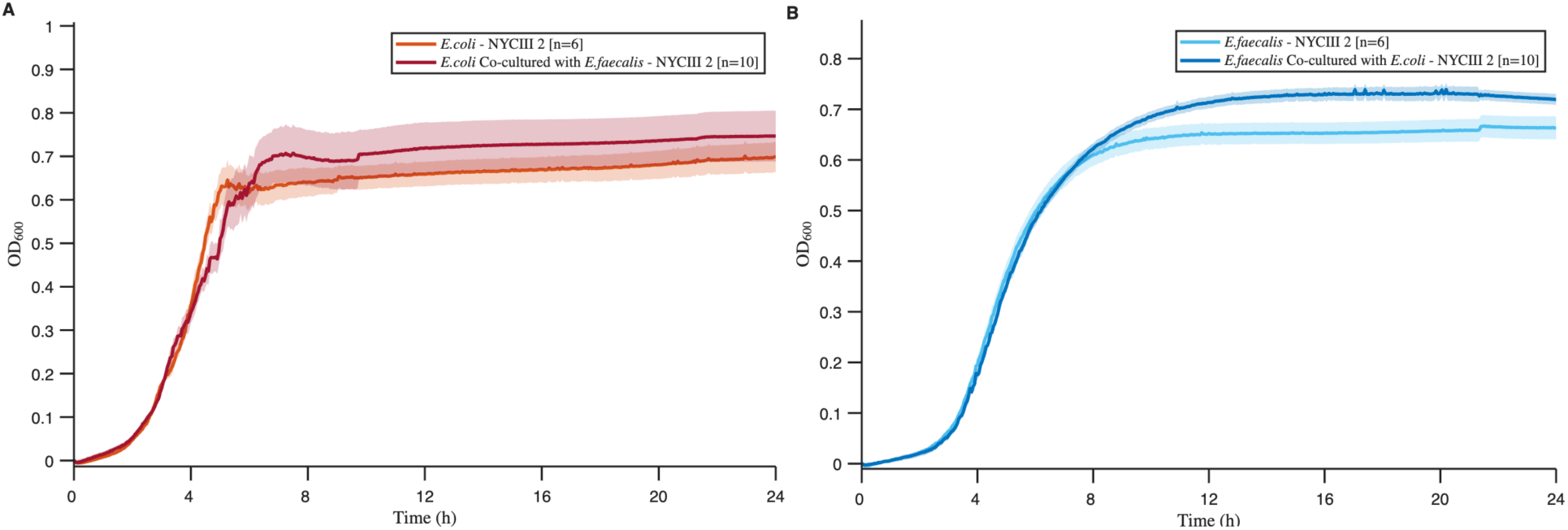
Bacterial growth curve in NYCIII medium of *E*. *coli* (ATCC 25922) (**A) and *E. faecalis* (ATCC 29212) in dual system.** Optical density at 600 nm (OD_600_) is shown on the y-axis; time (hours) on the x-axis. Lines represent mean values and shading standard error of the mean. The orange-colored line in panel A represents individual *E*. *coli* cultures whereas the light blue line in panel B represents individual *E*. *faecalis* cultures. The dark red line in panel A represents *E*. *coli* co-cultured with *E. faecalis* in opposing duet wells whereas the dark blue line in panel B represents *E*. *faecalis* co-cultured with *E. coli* in opposing duet wells.

**Fig. 11:**
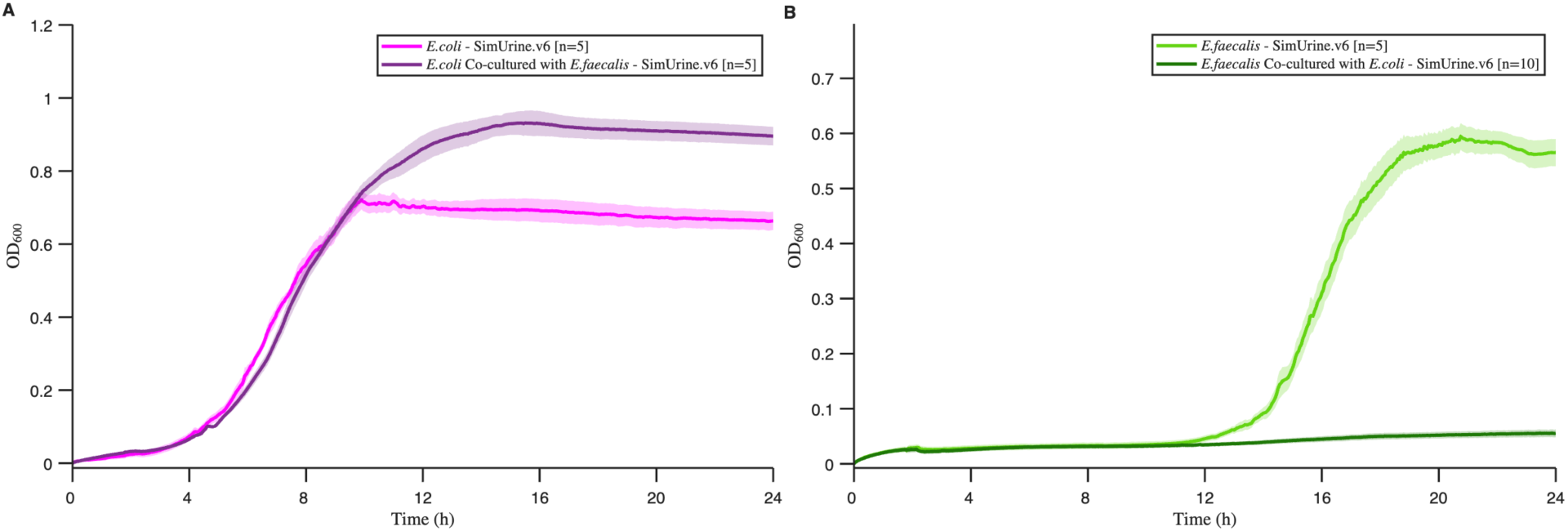
Bacterial growth curve in SimUrine.v6 medium of *E*. *coli* (ATCC 25922) (A) and *E. faecalis* (ATCC 29212) in dual system. Optical density at 600 nm (OD_600_) is shown on the y-axis; time (hours) on the x-axis. Lines represent mean values and shading standard error of the mean. The magenta line in panel A represents individual *E*. *coli* cultures whereas the light green line in panel B represents individual *E*. *faecalis* cultures. The plum-colored line in panel A represents *E*. *coli* co-cultured with *E.faecalis* whereas the dark green line in panel B represents *E*. *faecalis* co-cultured with *E. coli* in opposing duet wells.

This enhanced physiological relevance may lead to improved therapeutic decision-making and better understanding of urobiome dynamics in health and disease. The defined nature of SimUrine medium, and its physicochemical properties, makes it ideal for use in the study of the urobiome, using the duet system, microfluidic devices, organoid-on-a-chip devices, or simple flasks.

SimUrine cannot fully simulate several aspects of human urine. Certainly, it would be nearly impossible to account for the thousands of biomolecules that can be isolated in human urine, the majority of which are found in near trace amounts. However, even the most minute quantities of certain compounds may have physiological relevance when studying urobiome behaviors. SimUrine excludes hormones, xenobiotics, chromogens, cellular debris, mucoproteins, and macromolecules (DNA, collagen) due to instability, undefined composition, wide physiological ranges, or cost. However, SimUrine attempts to account for the vital contribution of glycosaminoglycans in the bladder environment towards bacterial survival with the inclusion of N-acetylglucosamine, L-threonine, and L-serine, all glycosaminoglycan degradation metabolites. It will be important to account for these missing variables when attempting to simulate bladder environments. Incorporating cell cultures such as urothelial cell lines or organoids may attempt to address some of these deficits.

Future applications of SimUrine include standardized urobiome research protocols, probiotic efficacy testing under physiologically relevant conditions, and development of personalized therapeutic approaches based on individual urine chemistry profiles. The medium’s defined composition also facilitates mechanistic studies of host-microbe interactions within the urinary tract ecosystem.

## Supplementary Data

**Supplementary Figure 1.**
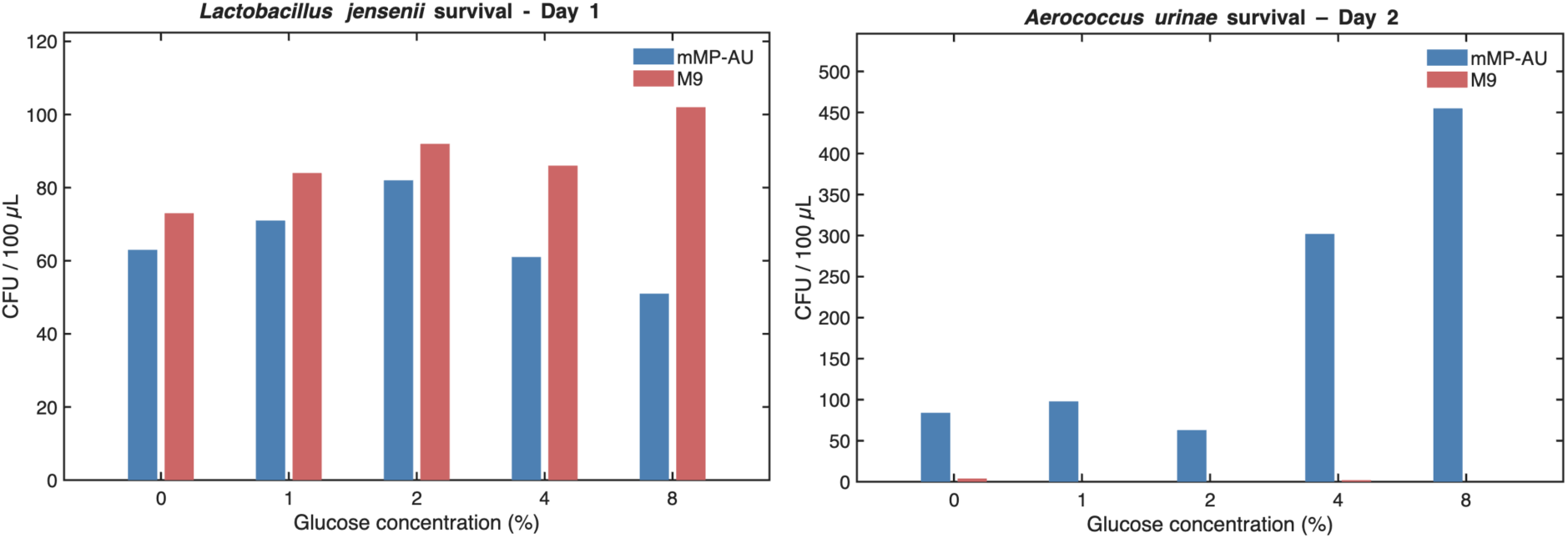
Effect of glucose supplementation on bacterial survival. (**A**) *Lactobacillus jensenii* (UMB8651) and (**B**) *Aerococcus urinae (*UMB5254) survival in mMP-AU or M9 salts supplemented with glucose. The y-axes display CFU/100 µL, and the x-axes represent glucose concentration.

**Supplementary Figure 2.**
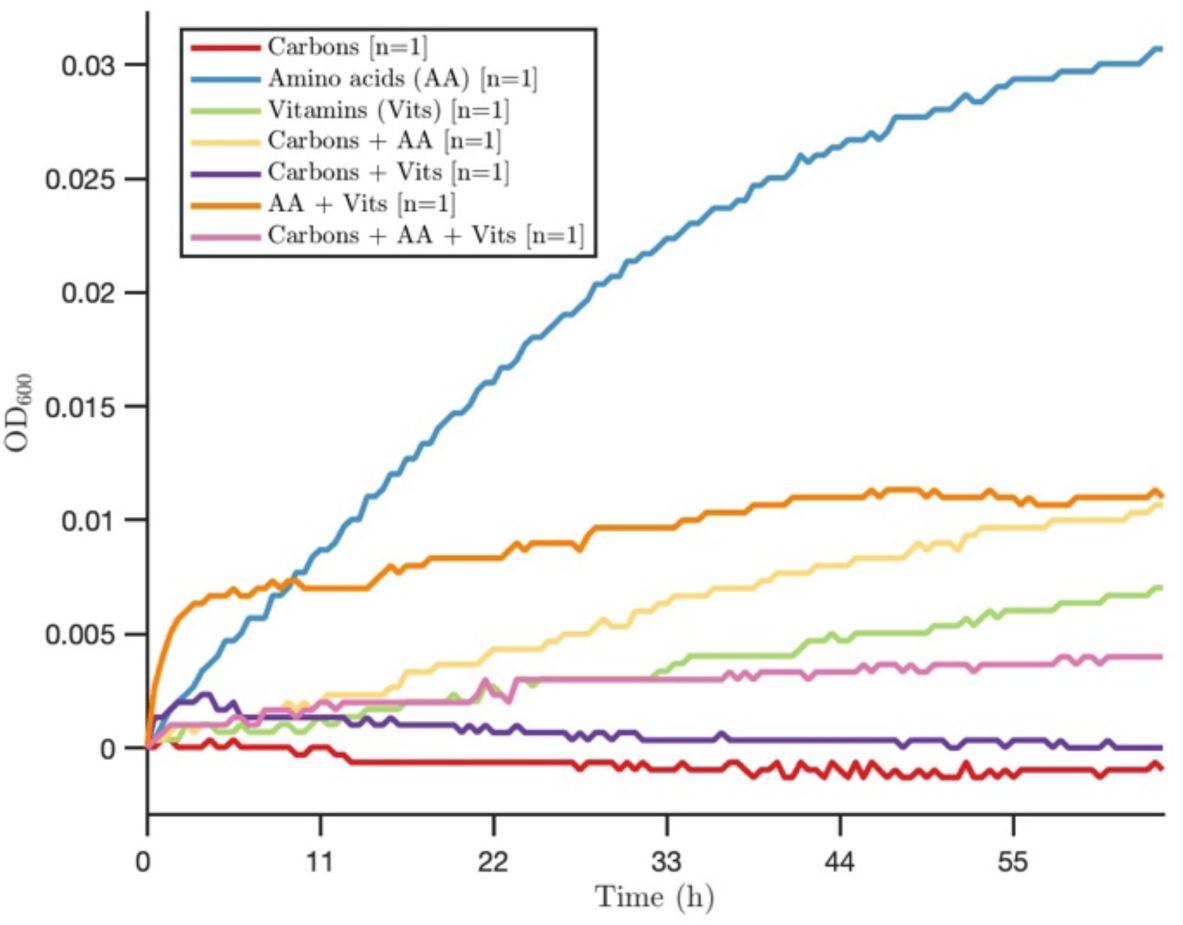
Effect of nutrient supplementation on *Lactobacillus jensenii* growth. Growth curves of *L. jensenii* (UMB8651) cultured in mMP-AU supplemented with carbon sources, amino acid sources, vitamins, or their combinations. Optical density at 600 nm (OD_600_) is shown on the y-axis; time (hours) on the x-axis

**Supplementary Figure 3: SimUrine.v1**. Left image shows the medium as it looked just out of the autoclave; right image, after 24 hours. Precipitate was formed within 24 hours.

**Supplementary Figure 4:**
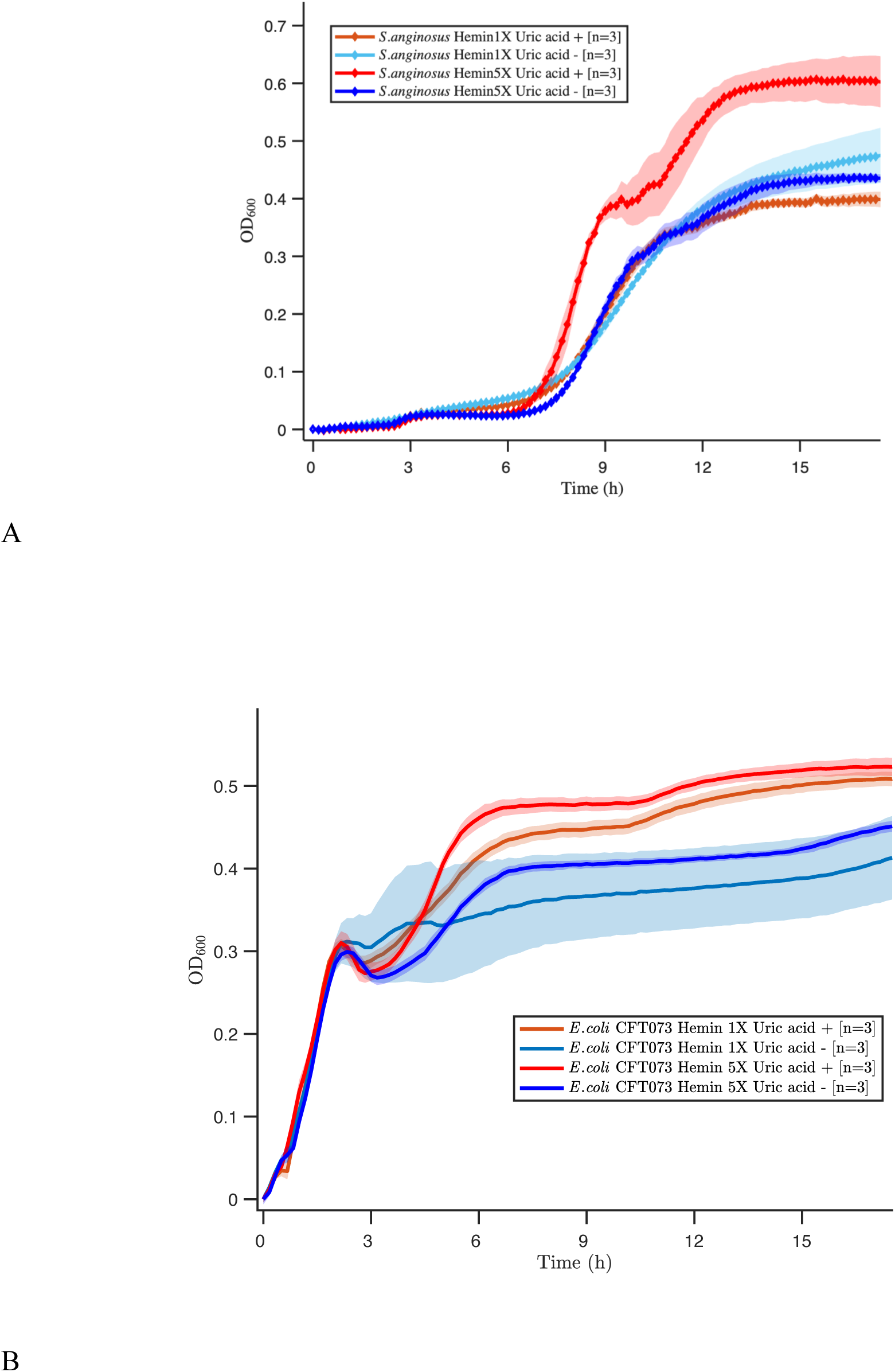
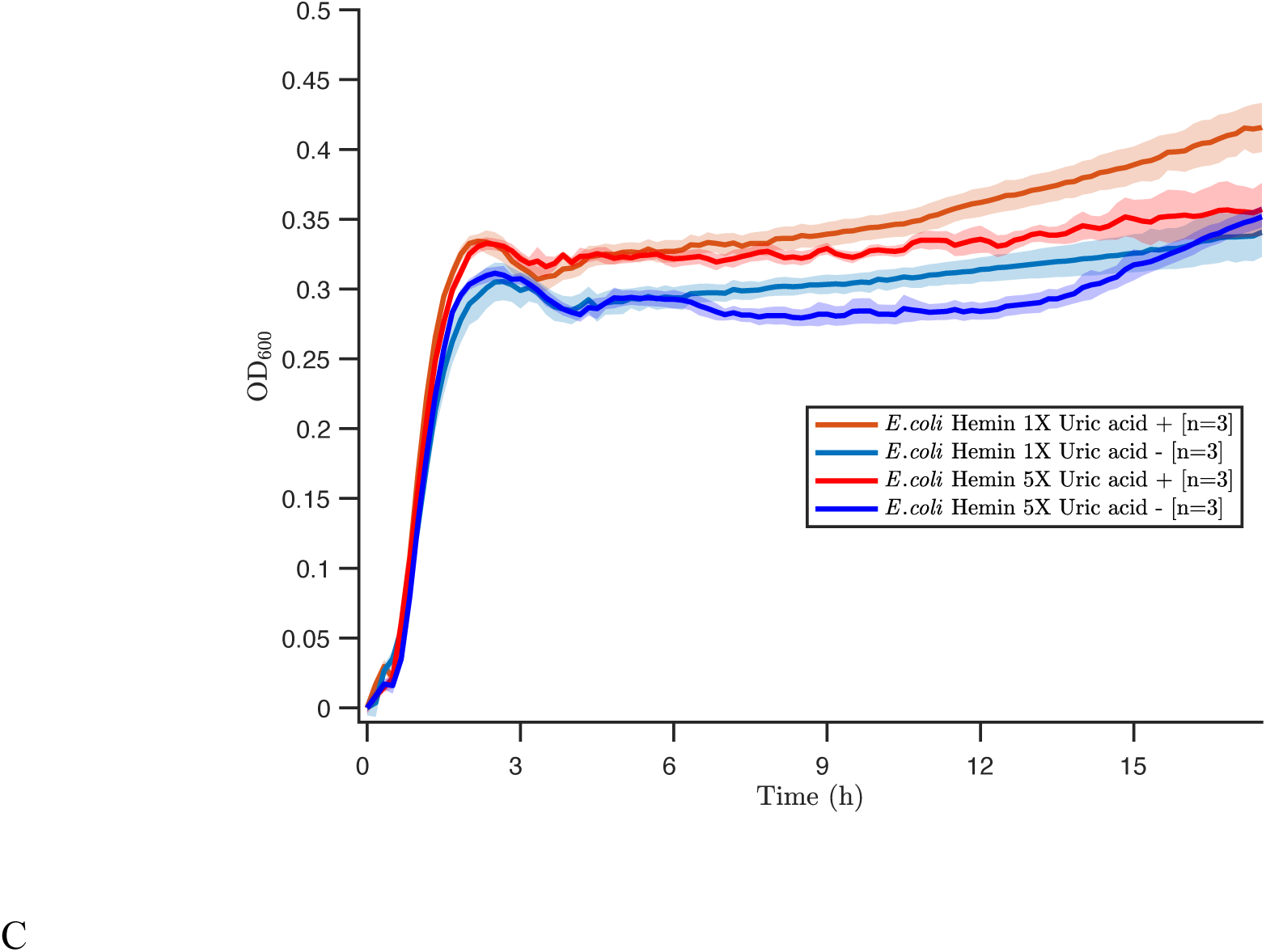
Cultures of *Streptococcus anginosus* (UBM 3192) (A), *E. coli* CFT073 (B) and *E. coli* (UBM3190) (C) in hemin-supplemented SimUrine.v3 in presence or absence of uric acid. Bold and light colors for high and low levels of hemin respectively, red tones for presence of uric acid and blue tones for absence. Optical density at 600 nm (OD600) is shown on the y-axis; time (hours) on the x-axis.

**Supplementary Table 1:** Strains used on this publication.

| Specie | Reference | Able to grow in SimUrine.V6 - 37°C |
| --- | --- | --- |
| *Escherichia coli* | UMB1180 | Not Tested |
| *Lactobacillus jensenii* | UBM8651 | Only in tubes, 5 % CO<sub>2</sub> |
| *Aerococcus urinae* | UMB5254 aka type strain<br>ATCC 51268 | Tubes and 96-well plates, 5% CO<sub>2</sub> |
| *Escherichia coli* | UMB3190, aka ATCC<br>10798 | Tubes and 96-well plates, 5% CO<sub>2</sub> |
| *Escherichia coli* (UPEC20) | NCBI BioSample database.<br>Accession<br>SAMN041053058.<br>National Library of<br>Medicine, National Center<br>for Biotechnology<br>Information<br>(RRID: SCR_004854). | Tubes and 96-well plates, 5% CO<sub>2</sub> |
| *Escherichia coli* | ATCC 25922 | Tubes and 96-well plates, 5% CO<sub>2</sub> |
| *Enterococcus faecalis* | ATCC 29212 | Tubes and 96-well plates, 5% CO<sub>2</sub> |
| *Klebsiella pneumoniae* | UBM9987 | Tubes and 96-well plates, 5% CO<sub>2</sub> |
| *Actinotignum schaalii* | UBM13319 | Only in glass tubes, 5 % CO<sub>2</sub> |
| *Streptococcus anginosus* | UBM8616 | Tubes and 96-well plates, 5% CO<sub>2</sub> |
| *Corynebacterium riegelii* | UBM12267 | Tubes and 96-well plates, 5% CO<sub>2</sub> |
| *Enterococcus faecalis* | UBM3193 | Tubes and 96-well plates, 5% CO<sub>2</sub> |
| *Enterococcus faecalis* | UBM7540 | Tubes and 96-well plates, 5% CO<sub>2</sub> |
| *Proteus mirabilis* (strain83) | Local strain (Croxall et al.,<br>2011) | Tubes and 96-well plates, 5% CO<sub>2</sub> |
| *Lactobacillus gasseri* | UMB4205 | Only in glass tubes, 5 % CO<sub>2</sub> |
| *Proteus mirabilis* | UBM7310 | Tubes and 96-well plates, 5% CO<sub>2</sub> |
| *Streptococcus anginosus* | UBM3192 | Tubes and 96-well plates, 5% CO<sub>2</sub> |
| *Lactobacillus gasseri* | UBM245 | Only in glass tubes, anaerobic |
| *Lactobacillus jensenii* | UBM387 | Only in glass tubes, anaerobic |
| *Prevotella buccalis* | UBM536 | Only in glass tubes, anaerobic |

### Supplementary Protocols 1

#### SimUrine.v6

**Protocol for 500mL of media 1X.**

Add 200 mL of H_2_O to a flask and dissolve the following reagents one by one stirring and keeping at 40°C.

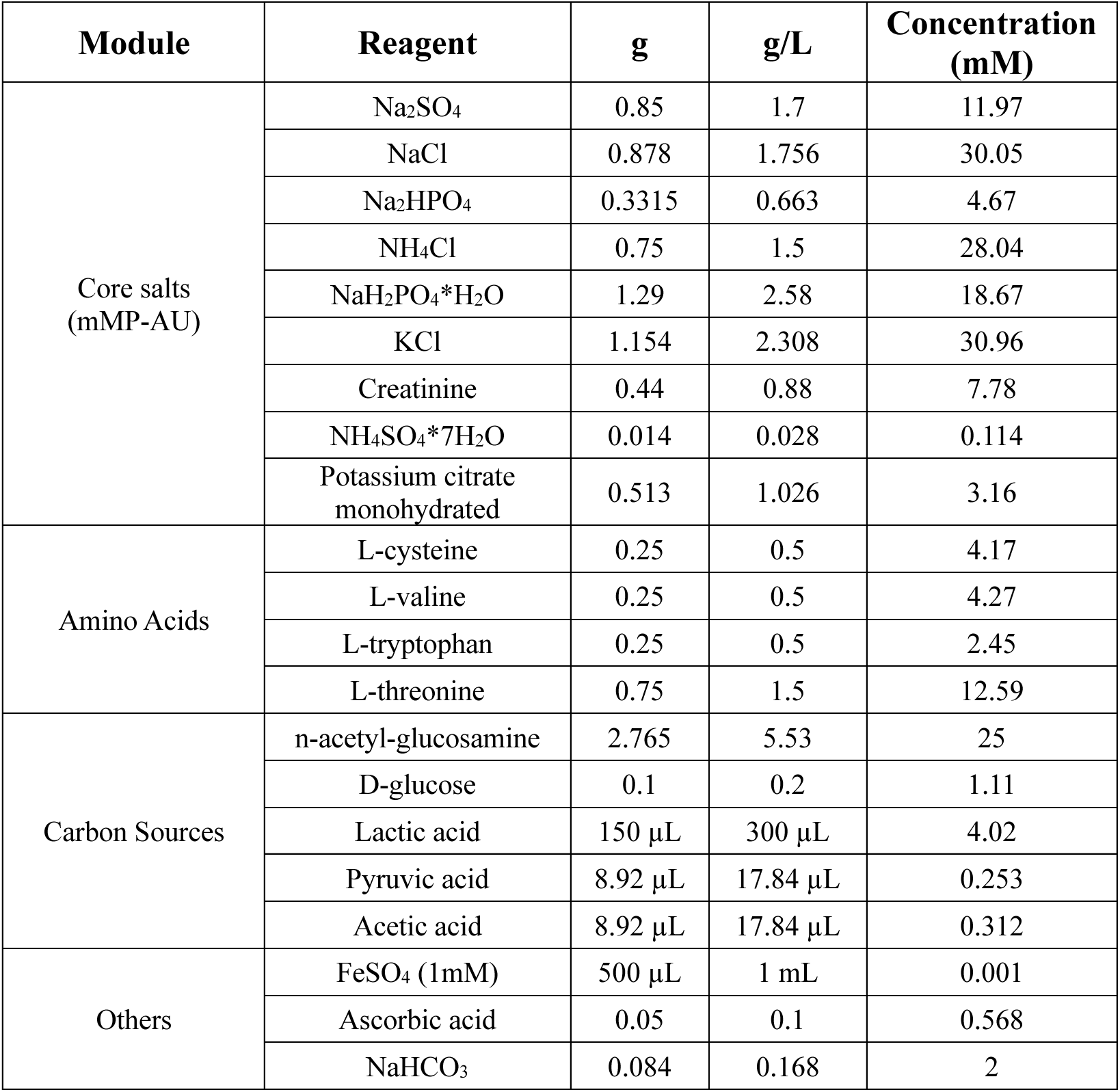

| Module | Reagent | g | g/L | Concentration (mM) |
| --- | --- | --- | --- | --- |
| Core salts (mMP-AU) | Na<sub>2</sub>SO<sub>4</sub> | 0.85 | 1.7 | 11.97 |
|  | NaCl | 0.878 | 1.756 | 30.05 |
|  | Na<sub>2</sub>HPO<sub>4</sub> | 0.3315 | 0.663 | 4.67 |
|  | NH<sub>4</sub>Cl | 0.75 | 1.5 | 28.04 |
|  | NaH<sub>2</sub>PO<sub>4</sub>*H<sub>2</sub>O | 1.29 | 2.58 | 18.67 |
|  | KCl | 1.154 | 2.308 | 30.96 |
|  | Creatinine | 0.44 | 0.88 | 7.78 |
|  | NH<sub>4</sub>SO<sub>4</sub>*7H<sub>2</sub>O | 0.014 | 0.028 | 0.114 |
|  | Potassium citrate monohydrated | 0.513 | 1.026 | 3.16 |
| Amino Acids | L-cysteine | 0.25 | 0.5 | 4.17 |
|  | L-valine | 0.25 | 0.5 | 4.27 |
|  | L-tryptophan | 0.25 | 0.5 | 2.45 |
|  | L-threonine | 0.75 | 1.5 | 12.59 |
| Carbon Sources | n-acetyl-glucosamine | 2.765 | 5.53 | 25 |
|  | D-glucose | 0.1 | 0.2 | 1.11 |
|  | Lactic acid | 150 µL | 300 µL | 4.02 |
|  | Pyruvic acid | 8.92 µL | 17.84 µL | 0.253 |
|  | Acetic acid | 8.92 µL | 17.84 µL | 0.312 |
| Others | FeSO<sub>4</sub> (1mM) | 500 µL | 1 mL | 0.001 |
|  | Ascorbic acid | 0.05 | 0.1 | 0.568 |
|  | NaHCO<sub>3</sub> | 0.084 | 0.168 | 2 |

While media is hot, add 500 µL of Tween-80.

Add 1.05g of MOPS

Add 7.5 g of Urea

Add:

a. 0.5 mL of solution of trace elements 1 (Acid)
b. 0.5 mL of solution of trace elements 2 (Basic)
c. 5 mL of vitamin mix 1
d. 5 mL of vitamin mix 2
e. 2 mL of solution of MgSO_4_ 108g/L (0.216 g).
f. 1 mL solution of NH_4_C_2_O_4_ 13.5 g/L (0.0135 g).
g. 100 µL of solution of **MnSO_4_** 0.8 g/L.
h. 5 mL of amino acids 100X solution.
i. 2500 µL of hemin solution 0.5 g/L (0.00125 g). **Check pH (it should be ∼ pH= 6, adapt it to pH=6 with HCl)**
j. 10 mL of uric acid solution 12.5 g/L (0.125 g). Add 0.25 g of L-serine **Check pH (it should be ∼ pH= 6, adapt it to pH=6 with HCl)**

**Complete volume to 500 mL with H_2_O. Filter using 0.2 µm filter. pH should stay stable.**

### SOLUTIONS

The following solutions must be prepared and sterilized before using them.

#### a) Trace elements (Acid). (Filter using a 0.2 µm filter. Protect from light, 4°C or RT)

| Reagent | mg | concentration |
| --- | --- | --- |
| CuCl<sub>2</sub> * 2H<sub>2</sub>O | 1.70 | 0.1 mM |
| H<sub>3</sub>BO<sub>3</sub> | 6.18 | 1 mM |
| CoCl<sub>2</sub>*6H<sub>2</sub>O | 11.89 | 0.5 mM |
| NiCl<sub>2</sub>*6H<sub>2</sub>O | 2.38 | 0.1 mM |
| ZnCl<sub>2</sub> | 6.82 | 0.5 mM |
| FeCl<sub>2</sub>*4H<sub>2</sub>O | 149.11 | 7.5 mM |

Dissolve in 100 mL of water and add 8.3 µL of HCl 37%.

#### b) Trace elements (Basic) (Filter using a 0.2 µm filter. 4°C or RT)

| Reagent | mg | concentration |
| --- | --- | --- |
| Na<sub>2</sub>SeO<sub>3</sub> | 1.73 | 0.1 mM |
| Na<sub>2</sub>WO<sub>4</sub>*2H<sub>2</sub>O | 3.3 | 0.1 mM |
| Na<sub>2</sub>MoO<sub>4</sub>*2H<sub>2</sub>O | 2.42 | 0.1 mM |

Dissolve in 100 mL of NaOH 10 mM

#### c) Vitamins 1 (Filter using a 0.2 µm filter. Protect from light, 4°C)

| Reagent | mg |
| --- | --- |
| Biotin | 2 |
| Nicotinamide | 20 |
| Thiamine | 20 |
| Riboflavin | 10 |

Dissolve in 100 mL of solution CaCl_2_ (15.25 g/L)

#### d) Vitamins 2 (Filter using a 0.2 µm filter. Protect from light, 4°C)

| Reagent | mg |
| --- | --- |
| p-aminobenzoic acid | 10 |
| Pantothenic acid | 10 |
| pyridoxamine | 50 |
| Cyanocobalamin | 10 |
| Folic acid | 10 |

Dissolve in 100 mL of solution CaCl_2_ (15.25 g/L)

#### e) MgSO_4_ solution (Autoclave or filter using a 0.2 µm filter. RT or 4°C. Indef. Storage)

MgSO₄·7H₂O 5.4 g in 50 mL of H_2_O

#### f) NH_4_C_2_O_4_ solution (Autoclave or filter. Indef. Storage)

NH_4_C_2_O_4_·H_2_O 1.35 g in 100 mL of H_2_O

#### g) MnSO4 solution (Indef. Storage)

MnSO_4_ 0.08 g in 100 mL of H_2_O.

#### h) Amino acids 100X solution (100 mL)

Add 40 mL of H_2_O and incorporate the water-soluble amino acids one by one. Dissolve individually the other amino acids in ∼ 5mL of HCl 1M and add to the bottle. Complete volume to 100 mL with H_2_O. To ensure solubility, keep solution at pH ∼1.

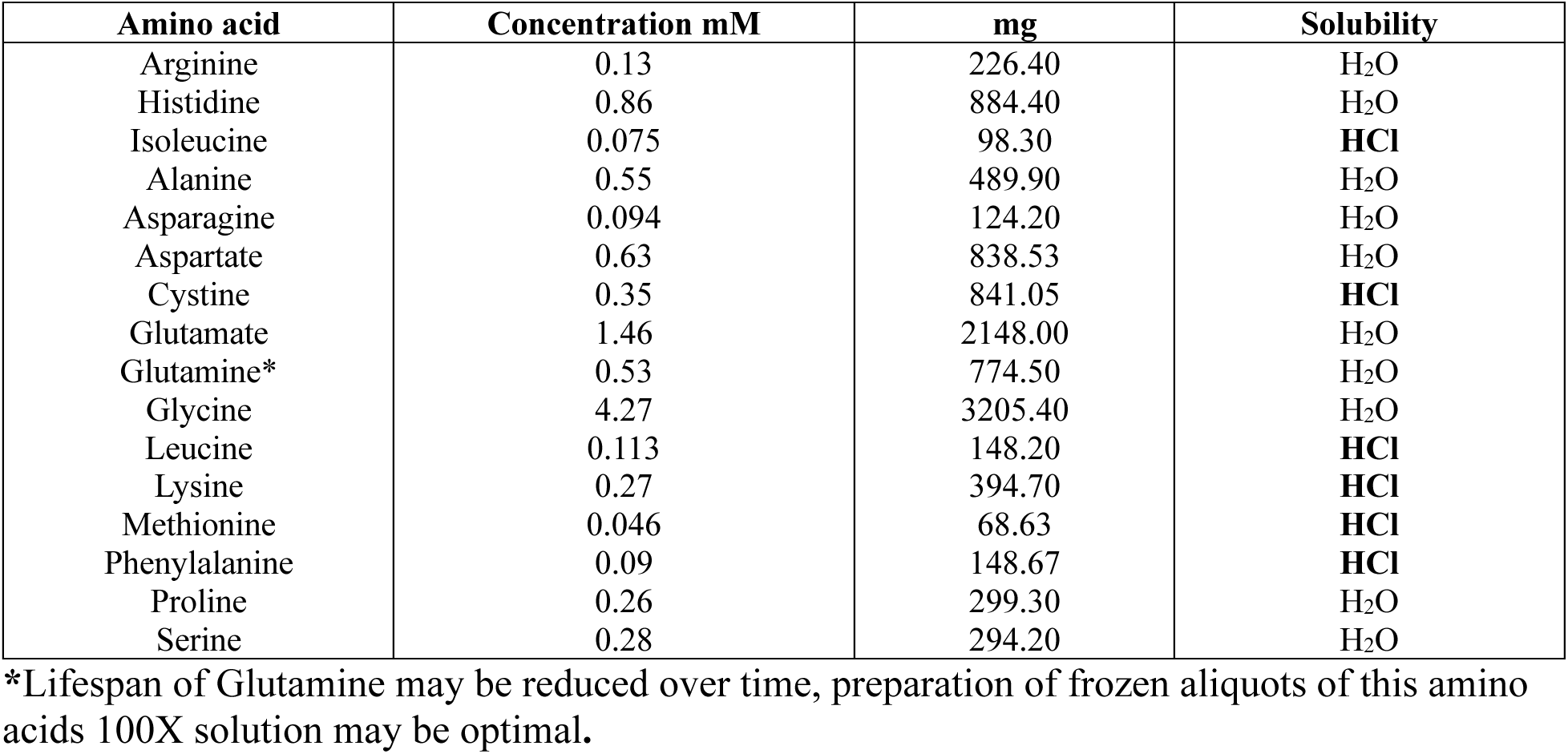

| Amino acid | Concentration mM | mg | Solubility |
| --- | --- | --- | --- |
| Arginine | 0.13 | 226.40 | H<sub>2</sub>O |
| Histidine | 0.86 | 884.40 | H<sub>2</sub>O |
| Isoleucine | 0.075 | 98.30 | HCl |
| Alanine | 0.55 | 489.90 | H<sub>2</sub>O |
| Asparagine | 0.094 | 124.20 | H<sub>2</sub>O |
| Aspartate | 0.63 | 838.53 | H<sub>2</sub>O |
| Cystine | 0.35 | 841.05 | HCl |
| Glutamate | 1.46 | 2148.00 | H<sub>2</sub>O |
| Glutamine* | 0.53 | 774.50 | H<sub>2</sub>O |
| Glycine | 4.27 | 3205.40 | H<sub>2</sub>O |
| Leucine | 0.113 | 148.20 | HCl |
| Lysine | 0.27 | 394.70 | HCl |
| Methionine | 0.046 | 68.63 | HCl |
| Phenylalanine | 0.09 | 148.67 | HCl |
| Proline | 0.26 | 299.30 | H<sub>2</sub>O |
| Serine | 0.28 | 294.20 | H<sub>2</sub>O |

\*Lifespan of Glutamine may be reduced over time, preparation of frozen aliquots of this amino acids 100X solution may be optimal.

#### i) Hemin solution (Autoclave, protect from light, 4°C)

Add 50 mg of Hemin in 1 mL of NaOH 1N, complete volume to 100 mL with H_2_O.

#### j) Uric acid solution (Autoclave, 4 days at 4°C)

0.625 g in 5 mL NaOH 10 M, dissolve, and complete to 50 mL with H_2_O.

**Supplementary Table 2:** List of reagents utilized in this work.

| Correlative | Reagents | Code | Company |
| --- | --- | --- | --- |
| 1 | Acetic acid | A6283-1L | Sigma-Aldrich |
| 2 | Alanine | 11442468 | Thermo scientific Chemicals |
| 3 | Arginine | A5006-100G | Sigma-Aldrich |
| 4 | Asparagine | A0884-25G | Sigma-Aldrich |
| 5 | Aspartate | A9256-100G | Sigma-Aldrich |
| 6 | Biotin | B4501-100MG | Sigma-Aldrich |
| 7 | CaCl<sub>2</sub> | C1016-100G | Sigma-Aldrich |
| 8 | Citric acid | C0759-100G | Sigma-Aldrich |
| 9 | CoCl<sub>2</sub>*6H<sub>2</sub>O | 202185-25G | Sigma-Aldrich |
| 10 | Creatinine | 228940500 | Thermo scientific Chemicals |
| 11 | CuCl<sub>2</sub> * 2H<sub>2</sub>O | 10588502 | Thermo scientific Chemicals |
| 12 | Cyanocobalamin | 1.2459201 | Sigma-Aldrich |
| 13 | Cystine | 11414814 | Thermo scientific Chemicals |
| 14 | FeCl<sub>2</sub>*4H<sub>2</sub>O | 10348150 | Thermo scientific Chemicals |
| 15 | Folic acid | F7876-10G | Sigma-Aldrich |
| 16 | Glutamate | 10502751 | Thermo scientific Chemicals |
| 17 | Glutamine | 11482277 | Thermo scientific Chemicals |
| 18 | Glycine | 1.00590.1000 | MERK |
| 19 | H<sub>3</sub>BO<sub>3</sub> | B0394-100G | Sigma-Aldrich |
| 20 | Hemin | 10499291 | Thermo scientific Chemicals |
| 21 | HEPES 1M | H3375-25G | Sigma-Aldrich |
| 22 | Histidine | H8000-5G | Sigma-Aldrich |
| 23 | Isoleucine | I2752-1G | Sigma-Aldrich |
| 24 | KCl | 1.04936.1000 | MERK |
| 25 | L-cysteine | A10435.18 | Thermo scientific Chemicals |
| 26 | L-serine | S4500-1G | Sigma-Aldrich |
| 27 | L-threonine | 1.0841101 | Sigma-Aldrich |
| 28 | Lactic acid | 10665302 | Thermo scientific Chemicals |
| 29 | Leucine | L8000-25G | Sigma-Aldrich |
| 30 | Lysine | L5501-10MG | Sigma-Aldrich |
| 31 | Methionine | M9625-25G | Sigma-Aldrich |
| 32 | MgSO<sub>4</sub>*7H<sub>2</sub>O | 447155000 | Thermo scientific Chemicals |
| 33 | MnSO<sub>4</sub>*H<sub>2</sub>O | 11478237 | Thermo scientific Chemicals |
| 34 | n-acetyl-glucosamine | 11438770 | Thermo scientific Chemicals |
| 35 | Na<sub>2</sub>HPO<sub>4</sub> | 1.06586.0500 | Sigma-Aldrich |
| 36 | Na<sub>2</sub>MoO<sub>4</sub>*2H<sub>2</sub>O | 10164830 | Thermo scientific Chemicals |
| 37 | Na<sub>2</sub>SO<sub>4</sub> | 1.28211.1001 | Sigma-Aldrich |
| 38 | Na<sub>2</sub>WO<sub>4</sub>*2H<sub>2</sub>O | 10117423 | Thermo scientific Chemicals |
| 39 | NaCl | 207790010 | ACROS |
| 40 | NaH<sub>2</sub>PO<sub>4</sub> | 1.06349.1000 | Sigma-Aldrich |
| 41 | NaOH | S5881-1K | Sigma-Aldrich |
| 42 | Na<sub>2</sub>SeO<sub>3</sub> | 214485-5G | Sigma-Aldrich |
| 43 | NH<sub>4</sub>C<sub>2</sub>O<sub>4</sub> | 10054013 | Thermo scientific Chemicals |
| 44 | NH<sub>4</sub>Cl | A4514-500G | Sigma-Aldrich |
| 45 | NH<sub>4</sub>SO<sub>4</sub>*7H<sub>2</sub>O | A5132-1K | Sigma-Aldrich |
| 46 | NiCl<sub>2</sub>*6H<sub>2</sub>O | 654507-5G | Sigma-Aldrich |
| 47 | Nicotinamide | 10447710 | Thermo scientific Chemicals |
| 48 | p-aminobenzoic acid | A9878-5G | Sigma-Aldrich |
| 49 | Pantothenic acid | 21210-5G-F | Sigma-Aldrich |
| 50 | Phenylalanine | 11468840 | Thermo scientific Chemicals |
| 51 | Proline | 11439512 | Thermo scientific Chemicals |
| 52 | pyridoxamine | 10785052 | Thermo scientific Chemicals |
| 53 | Pyruvic acid | 10562721 | Thermo scientific Chemicals |
| 54 | Riboflavin | 10593871 | Thermo scientific Chemicals |
| 55 | Serine | 11489113 | Thermo scientific Chemicals |
| 56 | Thiamine | 10477540 | Thermo scientific Chemicals |
| 57 | Tween80 | P4780-500ML | Sigma-Aldrich |
| 58 | Urea | U5378-1K | Sigma-Aldrich |
| 59 | Uric acid | U2625-100G | Sigma-Aldrich |
| 60 | ZnCl<sub>2</sub> | 208086-5G | Sigma-Aldrich |

### Supplementary Protocols 2

#### SimUrine.v1

**Protocol for 500 ml of media 1X.**

Add 200 ml of H_2_O to a bottle and add the following reagents one by one.

| Reagent | g | g/L |
| --- | --- | --- |
| Na<sub>2</sub>SO<sub>4</sub> | 0.85 | 1.7 |
| NaCl | 0.878 | 1.756 |
| Na<sub>2</sub>HPO<sub>4</sub> | 0.3315 | 0.663 |
| NH<sub>4</sub>Cl | 0.633 | 1.266 |
| NaH<sub>2</sub>PO<sub>4</sub> | 1.12 | 2.24 |
| KCl | 1.154 | 2.308 |
| n-acetyl-glucosamine | 2.765 | 5.53 |
| L-threonine | 0.75 | 1.5 |
| CaCl<sub>2</sub>•H<sub>2</sub>O | 0.122 | 0.244 |

Dissolve individually in 5 ml of HCl 1M and add to the media.

0.25 g L-cysteine

0.25 g L-serine

Add: Lactic acid 75 µL, Pyruvic acid 4.46 µL, Acetic acid 4.46 µL Dissolve individually in 20 ml of H_2_O and add to the media:

Creatinine 0.44 g, MgSO_4_*7H2O 0.54 g, NH_4_SO_4_*7H2O 0.014 g,

Potassium citrate 0.484 g

Add 60 ml of H_2_O to complete ∼350 ml.

Autoclave (adjust pH 6.5)

Add 10 ml of uric acid solution and 1 ml solution of NH_4_C_2_O_4_ (See Supplementary protocol 1).

Add 7.5 g of urea, adjust pH to 6.5, complete with sterile water, and filter.

#### SimUrine.v2

**Protocol for 500 ml of media 1X.**

Add 200 ml of H_2_O to a bottle and add the following reagents one by one.

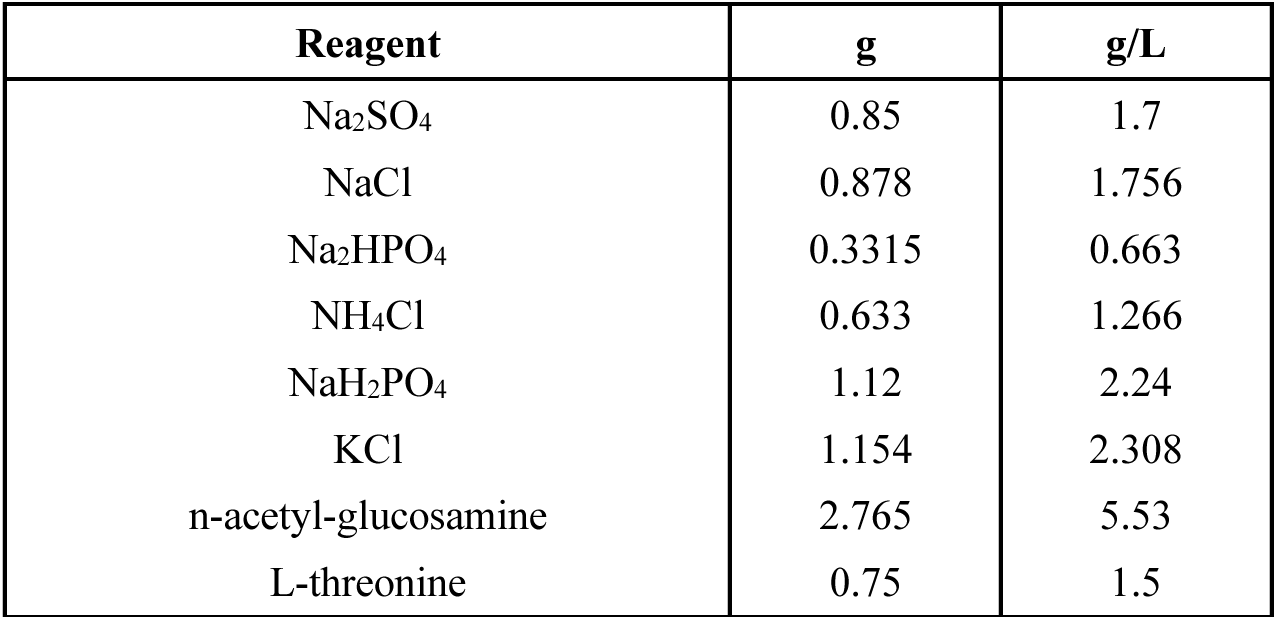

Dissolve individually in 5 ml of HCl 1M and add to the media.

0.25 g L-cysteine

0.25 g L-serine

Add: Lactic acid 75 µL, Pyruvic acid 4.46 µL, Acetic acid 4.46 µL

Dissolve individually in 20 ml of H_2_O and add to the media:

Creatinine 0.44 g, MgSO_4_*7H2O 0.54 g, NH_4_SO_4_*7H2O 0.014 g,

Potassium citrate 0.484 g

Add 60 ml of H_2_O to complete ∼350 ml.

Autoclave (adjust pH 6.5)

Add 5 ml of vitamin mix 1, 5 ml of vitamin mix 2*, 1 ml solution of NH_4_C_2_O_4_, 0.5 ml of solution of trace elements 1 (Acid), 0.5 ml of solution of trace elements 2 (Basic). (See **Supplementary Protocol 1**).

Add 10 ml of uric acid solution (12.5 g/L).

Add 7.5 g of urea, adjust pH to 6.5, complete with sterile water, and filter.

#### SimUrine.v3

**Protocol for 500 ml of media 1X.**

Add 200 ml of H_2_O to a bottle and add the following reagents one by one.

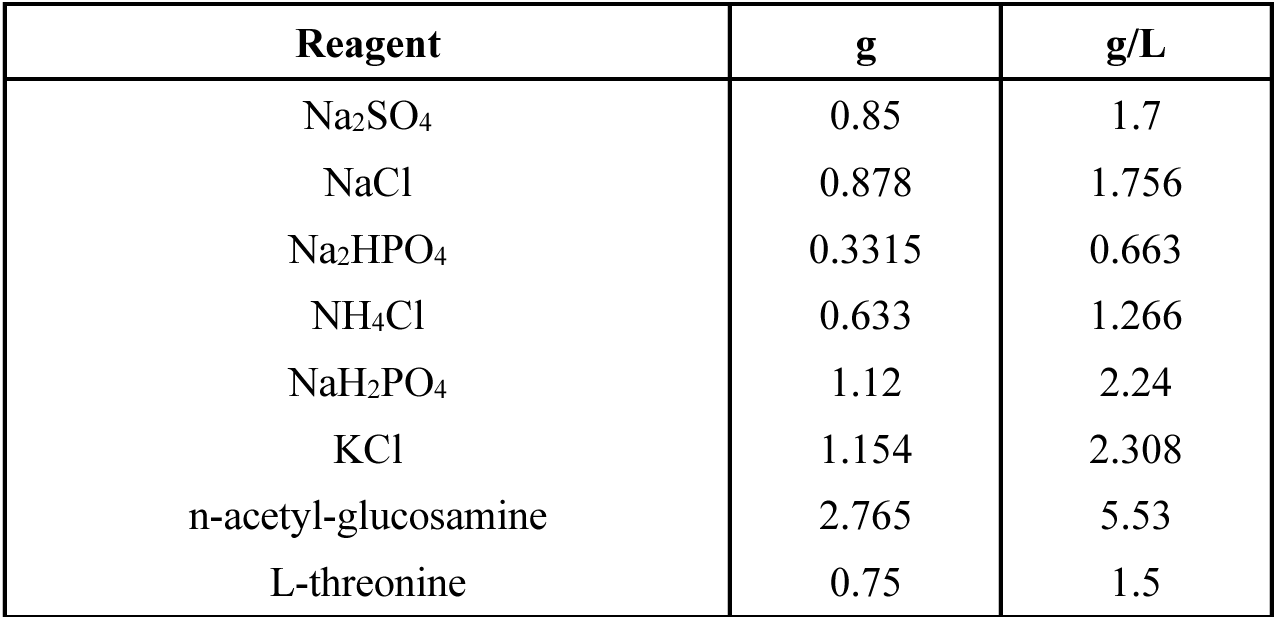

Dissolve individually in 5 ml of HCl 1M and add to the media.

0.25 g L-cysteine

0.25 g L-serine

Add: Lactic acid 75 µL, Pyruvic acid 4.46 µL, Acetic acid 4.46 µL Dissolve individually in 20 ml of H_2_O and add to the media:

Creatinine 0.44 g, MgSO_4_*7H2O 0.54 g, NH_4_SO_4_*7H2O 0.014 g

Potassium citrate 0.484 g

Add 60 ml of H_2_O to complete ∼350 ml.

Autoclave (adjust pH 6.5)

Add 5 ml of vitamin mix 1, 5 ml of vitamin mix 2*, 1 ml solution of NH_4_C_2_O_4_, 0.5 ml of solution of trace elements 1 (Acid), 0.5 ml of solution of trace elements 2 (Basic), 2500 µL of hemin solution (See Supplementary protocol 1).

Add 10 ml of uric acid solution (12.5 g/L).

Add 7.5 g of urea, adjust pH to 6.5, complete with sterile water, and filter.

#### SimUrine.v4

**Protocol for 500 ml of media 1X.**

Add 200 ml of H_2_O to a bottle and add the following reagents one by one.

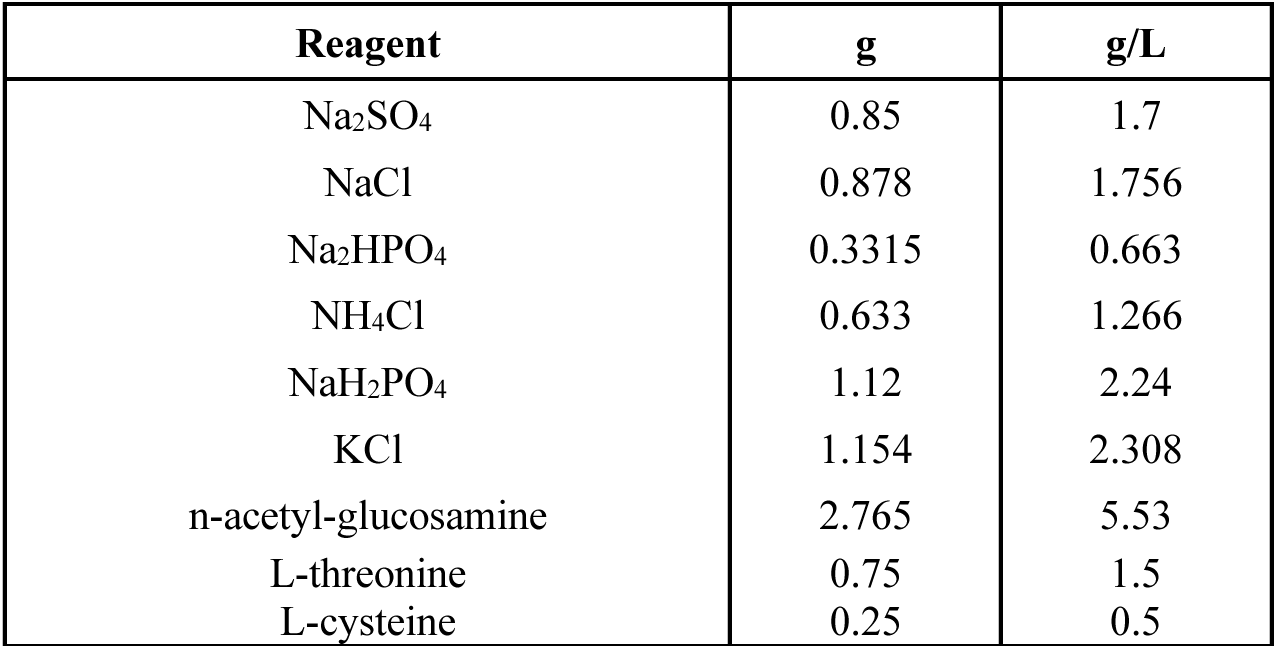

Dissolve individually in 5 ml of HCl 1M and add to the media.

0.25 g L-cysteine

0.25 g L-serine

Add: Lactic acid 75 µL, Pyruvic acid 4.46 µL, Acetic acid 4.46 µL Dissolve individually in 20 ml of H_2_O and add to the media:

Creatinine 0.44 g, NH_4_SO_4_*7H2O 0.014 g

Potassium citrate 0.484 g

Add 60 ml of H_2_O to complete ∼350 ml.

Autoclave (adjust pH 6.5)

**After autoclaving, add:**

(Here and while media is hot, add Tween-80 1 ml per liter for *Lactobacillus* growth.)

a. 0.5 ml of solution of trace elements 1 (Acid)
b. 0.5 ml of solution of trace elements 2 (Basic)
c. 5 ml of vitamin mix 1
d. 5 ml of vitamin mix 2
e. 1 ml of solution of MgSO_4_
f. 1 ml solution of NH_4_C_2_O_4_.
g. 100 µL of solution of MnSO_4_.

Adjust pH to 6.5 (∼100µL of NaOH 10M)

**(Check/adjust pH (6.5) and add 5 ml of HEPES 1M)**

Add 7.5 g of **urea** and filter.

**After filtering**, add:

h) 5 ml of amino acids 10X solution.

i) 2500 µL of hemin solution. (check pH, add ∼500 µL HCl 1M. Final pH 6.5)

j) 10 ml of uric acid solution.

**Complete volume to 500 ml with autoclaved H_2_O. pH should stay stable.**

#### SimUrine.v5

**Protocol for 500 ml of media 1X.**

Add 200 ml of H_2_O to a bottle and add the following reagents one by one.

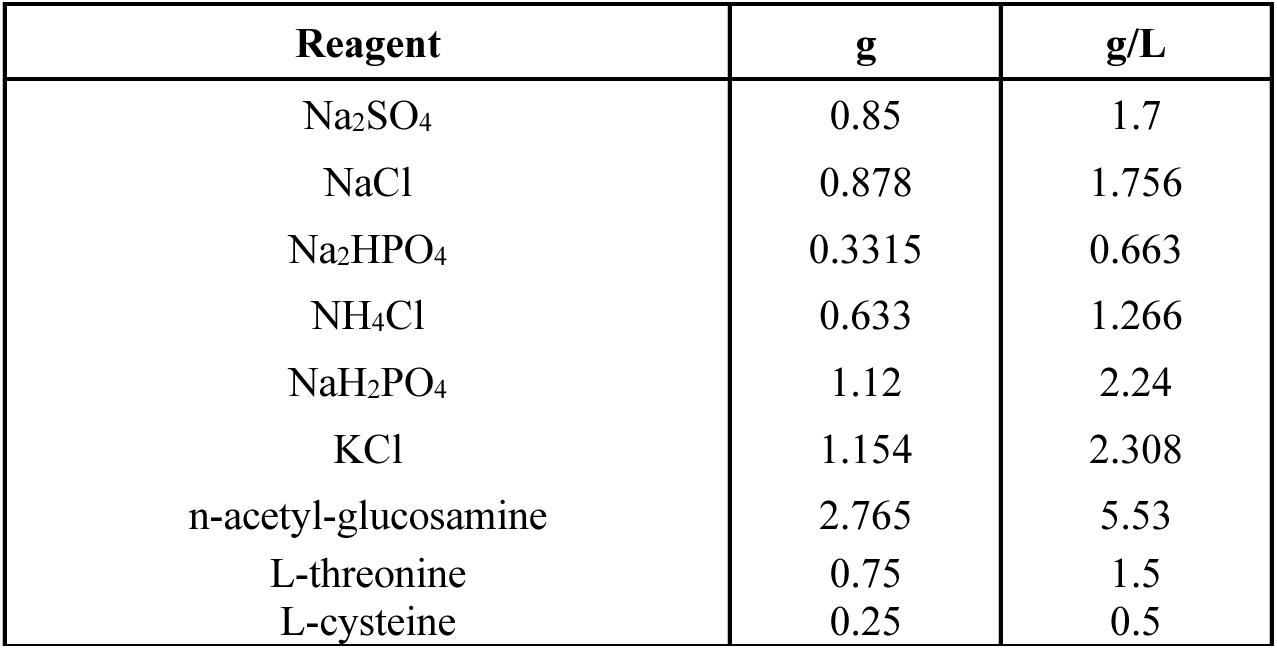

Dissolve individually in 5 ml of HCl 1M and add to the media.

0.25 g L-cysteine

0.25 g L-serine

Add: Lactic acid 75 µL, Pyruvic acid 4.46 µL, Acetic acid 4.46 µL Dissolve individually in 20 ml of H_2_O and add to the media:

Creatinine 0.44 g, NH_4_SO_4_*7H2O 0.014 g

Potassium citrate 0.484 g

Add 60 ml of H_2_O to complete ∼350 ml.

Autoclave (adjust pH 6.5)

**After autoclaving, add:**

(Here and while media is hot, add Tween-80 1 ml per liter for *Lactobacillus* growth.)

a. 0.5 ml of solution of trace elements 1 (Acid)
b. 0.5 ml of solution of trace elements 2 (Basic)
c. 5 ml of vitamin mix 1
d. 5 ml of vitamin mix 2
e. 1 ml of solution of MgSO_4_
f. 1 ml solution of NH_4_C_2_O_4_.
g. 100 µL of solution of MnSO_4_.

Adjust pH to 6 (∼100µL of NaOH 10M)

**(Check/adjust pH (6) and add MOPS 2.1 g/L)**

Add **2.5 g** of **urea** and filter.

**After filtering**, add:

k) 5 ml of amino acids 100X solution.

l) 2500 µL of hemin solution. (check pH, add ∼500 µL HCl 1M. Final pH 6.5)

m) 10 ml of uric acid solution.

**Complete volume to 500 ml with autoclaved H_2_O. pH should stay stable.**

## Funding

NWO ENW-XL OCENW.XL21.XL21.088

## Disclosures

AJW discloses compensated membership on the following scientific advisory boards: Astek, Cerillo, Pathnostics, and Urobiome Therapeutics. All other authors have no disclosures.

## Acknowledgments

We wish to thank Jacob Hogins and Larry Reitzer (UT-Dallas) for their suggested modifications to the SimUrine formulation, to Kathrin Tomasek for performing the osmolarity tests of our media, to Ayla Kwant for the conductivity and turbidity testing, and to Peter Dijkstra and Julien Es Sayed for their help testing viscosity and other physicochemical parameters.

## Author Contributions

Conceptualization: AJW

Funding Acquisition: MdeV, AJW

Project Administration: MdeV, AJW

Methodology: PGM, BIC

Data Acquisition: PGM, BIC, MV, MHK

Analysis: PGM, BIC

Writing Original Draft: PGM

Review and Edit: PGM, BIC, MV, MHK, MdeV, AJW

## Notes

### Summary of Updates

Manuscript has been reviewed and resubmitted to correct typographical error in figure 8A and included missing figure 9B.

